# Rv3839-Rv3840 links the endogenous heme biosynthesis pathway with *Mycobacterium tuberculosis* adaptation to nitric oxide and iron limitation stress

**DOI:** 10.64898/2026.02.17.706279

**Authors:** Natalia F. Quirk, Kate N. Gregory, Yasu S. Morita, Shumin Tan

**Affiliations:** Department of Molecular Biology and Microbiology, Tufts University School of Medicine, Boston, Massachusetts, USA; Graduate Program in Molecular Microbiology, Graduate School of Biomedical Sciences, Tufts University, Boston, Massachusetts, USA; Department of Microbiology, University of Massachusetts Amherst, Amherst, Massachusetts, USA; Molecular and Cellular Biology Graduate Program, University of Massachusetts Amherst, Amherst, Massachusetts, USA

## Abstract

During infection, *Mycobacterium tuberculosis* (Mtb) encounters multiple environmental stressors, including nitric oxide (NO) and iron limitation, and an ability to mount an integrated response is essential for the bacterium’s adaptation and continued survival. Iron-containing prosthetic groups in key enzymes are critical for Mtb sensing and detoxification of NO, and there is significant overlap between NO- and low iron-responsive genes. However, how Mtb adapts to these two stressors concurrently is largely unknown. Here, we find that exposure to NO globally augments expression of low iron-responsive genes and vice versa, with a two gene operon, *rv3839-rv3840,* among the most highly upregulated. Deletion of *rv3839-rv3840* resulted in increased growth under prolonged iron limitation and early exit of Mtb from an adaptive state of growth arrest induced upon exposure to NO/low iron. Δ*rv3839-rv3840* Mtb exhibited an elongated cell morphology compared to wild type Mtb in NO/low iron conditions, indicating effects of this operon on cell growth and division under stress conditions, with Rv3839 as the key driver of this phenotype. Coproporphyrin III tetramethyl ester (TMC), a modified precursor molecule in the endogenous Mtb heme biosynthesis pathway, was found to accumulate in Δ*rv3839-rv3840* Mtb under iron limiting conditions. Further, intrabacterial heme levels were increased in Δ*rv3839-rv3840* Mtb under NO stress and iron limitation. Together, these findings reveal Rv3839-Rv3840 as proteins involved in the downregulation of heme biosynthesis under NO stress and iron limitation, and highlight the link between Mtb growth control in response to NO/low iron and endogenous heme biosynthesis.

**IMPORTANCE:** Slowed growth is a physiologic adaptation to key environmental cues important for survival of *Mycobacterium tuberculosis* (Mtb) in the host. Nitric oxide (NO) is one such signal, but while regulation of NO response by the DosRS(T) two-component system is well-studied, NO stress also provokes a broad transcriptional response outside of DosRS(T) regulation that overlaps with the transcriptional response to iron limitation. Here, we show that Rv3839-Rv3840 contribute to Mtb maintenance of NO and low iron-induced growth arrest and find that this inability to maintain growth arrest is connected to dysregulation of the endogenous heme biosynthetic pathway. Little is known about regulation of endogenous heme biosynthesis in Mtb and its role in Mtb survival under stress conditions, and our results reveal a previously unknown interplay between NO and iron limitation response regulation and heme homeostasis.

## INTRODUCTION

Tuberculosis remains a particularly difficult disease to treat in part because of the ability of *Mycobacterium tuberculosis* (Mtb) to persist in the face of host defense mechanisms (1). Over the course of infection, Mtb encounters a range of environmental signals including nitric oxide (NO), hypoxia, and iron limitation. The environmental cues Mtb encounters varies depending on factors such as the infection stage (e.g. pre- or post-adaptive immune response onset) and its location within the host (e.g. necrotic core versus macrophage-rich cuff of a lesion) (2–4). Sensing and integrating response to these various signals is critical for Mtb adaptation to its local environment, and its ability to do so is vital for the bacterium’s continued survival *in vivo*, as evidenced by the significant attenuation in host colonization of Mtb lacking key regulators (5–7). However, while several key regulators have been identified (6–11), much remains unknown as to how Mtb integrates its response to different signals in its environment.

A key adaptive output of Mtb in response to environmental stress is alteration of its growth status. For example, Mtb growth is slowed at acidic pH (12, 13), and the bacteria also enter an adaptive state of growth arrest upon extended exposure to NO or hypoxia (11, 14, 15). Strikingly, a point mutation in a single two-component system (TCS) response regulator that resulted in faster growth of Mtb *in vitro* and failure to enter into a state of growth arrest upon extended NO exposure conversely resulted in attenuation for *in vivo* host colonization (8). This supports that slowed growth under certain environmental pressures is advantageous for Mtb during host colonization. NO stress, along with hypoxia, provokes a transcriptional response regulated by the DosRS(T) TCS in a set of 48 genes known as the “dormancy regulon” (6, 10, 11), and the two environmental signals have thus most often been studied together and in the context of DosR regulation. However, Mtb additionally encounters NO independently of hypoxia as part of the host adaptive immune response (16–19). RNA sequencing (RNAseq) data indicate that as many as 100 genes are significantly differentially expressed in response to NO that are not responsive to hypoxia (8, 20). Therefore, the response of Mtb to NO and hypoxia is differentiated, despite traditionally being studied in the context of their co-regulation. Further, DosR regulates only a subset (48) of the genes differentially expressed in response to NO (6, 8, 10, 11), indicating that our understanding of the regulatory network underlying the NO response of Mtb and how it may integrate with other aspects of Mtb cell biology is incomplete.

Notably, published transcriptional data show a significant overlap (46 genes) in Mtb transcriptional response to NO and iron limitation (8, 21, 22). During infection, host cells sequester free iron to starve pathogens of iron, an essential cofactor for many cellular processes (23). However, Mtb has an arsenal of iron binding, transport, and storage mechanisms, and can additionally utilize host heme as an iron source (24, 25). These strategies together allow Mtb to successfully compete with the host for iron (23, 26) and are intertwined with the Mtb response to NO stress. For example, expression of much of the iron response machinery is controlled by the essential iron-dependent transcriptional regulator IdeR (Rv2711) (21, 27–29), which has been shown to be required for resistance to reactive nitrogen intermediates (28). NO exposure degrades iron-sulfur (Fe-S) clusters in key enzymes, and Mtb must therefore upregulate iron acquisition and Fe-S cluster biogenesis as part of its defense against NO stress (30). Additionally, iron is required for heme production, which is involved in both the sensing and detoxification of NO stress. NO is directly sensed by the DosRS(T) TCS via heme prosthetic groups (31), and truncated hemoglobin N (HbN) detoxifies NO through its potent dioxygenase activity (32–34). Together, these findings support the existence of critical links between the response of Mtb to simultaneous NO stress and iron limitation, two environmental cues that would be encountered together during infection.

Here, we find that the Rv3839*-*Rv3840 proteins, which are encoded together in an operon and highly expressed upon exposure to both NO stress and iron limitation, limit Mtb growth under iron-limiting conditions and contribute to the maintenance of NO and iron stress-induced growth arrest. Iron limitation resulted in elongated Mtb cells, and while wild type (WT) Mtb lost this phenotype when acute NO stress was introduced, Δ*rv3839-rv3840* Mtb continued to exhibit an elongated morphology. Further, deletion of *rv3839-rv3840* resulted in accumulation of coproporphyrin III tetramethyl ester (TMC) under iron-limiting stress and increased intracellular heme. Together, these data reveal a previously unknown aspect of NO and iron limitation response regulation in Mtb and support an intrinsic interplay between Mtb adaptation to NO stress, iron limitation, and heme homeostasis.

## RESULTS

### Transcriptional response of Mtb to NO stress is augmented in the presence of iron limitation and vice versa

To first understand the intersection between Mtb response to iron limitation and NO stress, we sought to define the global transcriptional response to simultaneous NO stress and iron limitation via RNA sequencing (RNAseq) analysis with Mtb exposed to conditions of iron limitation, NO stress, or both. Notably, the RNAseq dataset showed augmentation in the induction of 94 out of 135 NO-responsive genes when NO stress is compounded with iron limitation (genes differentially expressed log_2_-fold change ≥1 in the NO condition; log_2_-fold change ≥0.6 between NO + low iron and NO conditions; p<0.05, FDR<0.01 in both sets) (Fig. 1A and Table S1). Similarly, induction of 75 out 142 low iron-responsive genes was augmented in the simultaneous presence of NO stress (genes differentially expressed log_2_-fold change ≥1 in the low iron condition; log_2_-fold change ≥0.6 between NO + low iron and low iron conditions; p<0.05, FDR<0.01 in both sets) (Fig. 1B and Table S2). Overall, 780 genes were differentially expressed (log_2_-fold change ≥1) under simultaneous NO stress and iron limitation, with 151 genes specifically differentially expressed in the dual condition, but not in either single condition alone (Tables S1-S3). qRT-PCR analysis on genes exhibiting the largest expression changes under simultaneous NO stress and iron limitation versus the single conditions validated the results observed in the RNAseq dataset (Fig. 1C). These data support the concept that NO stress exacerbates the iron limitation experienced by Mtb and vice versa.

**Fig. 1.**
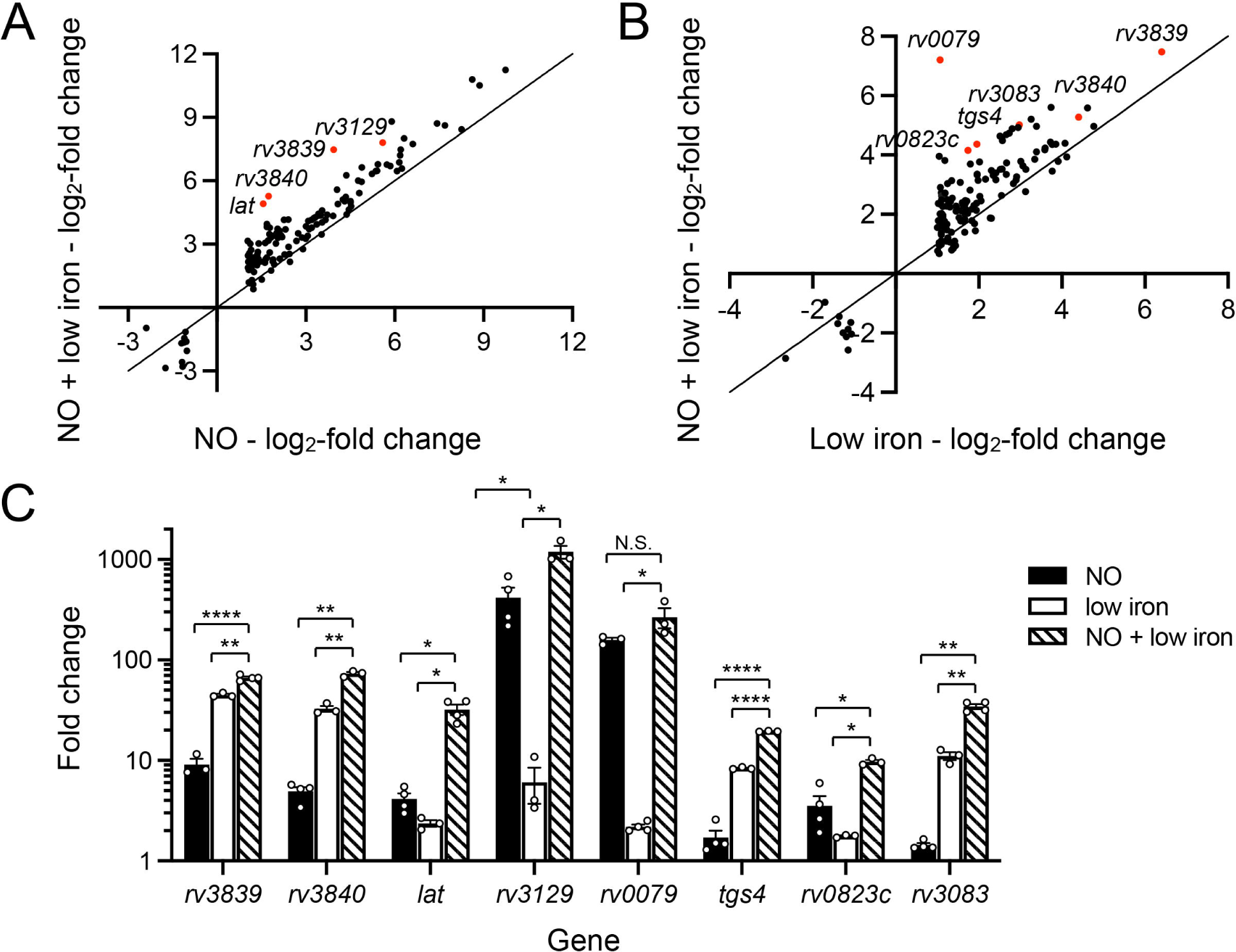
Mtb transcriptional response to NO stress is globally augmented in the presence of iron limitation and vice versa. Log-phase WT Mtb was exposed for 4 hours to: (i) iron-depleted minimal media + 150 µM FeNO_3_ (control), (ii) iron-depleted minimal media + 150 µM FeNO_3_ + 100 µM DETA NONOate (NO), (iii) iron-depleted minimal media + 100 µM 2’2’-dipyridyl (low iron), or (iv) iron-depleted minimal media + 100 µM 2’2’-dipyridyl + 100 µM DETA NONOate (NO + low iron), before RNA was extracted for RNAseq (A and B) or qRT-PCR (C). For RNAseq data, log_2_-fold change compares gene expression in each indicated condition to the control condition. Genes marked in red were the most highly augmented genes in NO + low iron compared to NO (A) or low iron (B) respectively (p<0.05, FDR<0.01 in both sets, with log_2_-fold change ≥1 in NO or low iron respectively). For qRT-PCR data, fold change is as compared to the control condition. *sigA* was used as the control gene and data are shown as means ± SEM from 3-4 experiments. p-values were obtained with an unpaired t-test with Welch’s correction and Holm-Sidak multiple comparisons. N.S. not significant, * p<0.05, ** p<0.01, **** p<0.0001.

### Rv3839-Rv3840 is important for maintenance of NO and iron stress-induced growth arrest

From the RNAseq data, the *rv3839-rv3840* operon stood out as it was among the most highly increased in expression in the dual NO + iron limitation condition compared to NO stress alone (Fig. 1A and C) and robustly upregulated under iron limitation (Fig. 1B). *rv3840*, encoding a gene annotated as a transcription factor (although it lacks a DNA-binding domain), is downstream of *rv3839*, which encodes a conserved hypothetical protein containing a domain of unknown function (DUF2470) found in heme utilization proteins (35, 36). Of note, *rv3839-rv3840* expression is induced by iron-limiting conditions in an IdeR-dependent manner (21), and *bfrB* (*rv3841*), encoding bacterioferritin B, an iron storage protein, is located immediately downstream of this operon. However, the function of Rv3839 and Rv3840 in Mtb is unknown. To determine if Rv3839-Rv3840 plays a role in regulating the Mtb iron limitation response, we generated a Δ*rv3839-rv3840* Mtb mutant and tested its growth under iron-limiting conditions. In standard 7H9 rich medium, Δ*rv3839-rv3840* Mtb grew indistinguishably from WT Mtb (Fig. 2A). In contrast, under iron-limiting conditions, Δ*rv3839-rv3840* Mtb exhibited increased growth compared to WT Mtb, which was most clearly observed with continued sub-culturing in iron-limiting conditions (Fig. 2A and B). Complementation of the *rv3839-rv3840* operon (*rv3839-rv3840\**) partially returned growth to WT levels (Fig. 2A and B). These data suggest that Rv3839*-*Rv3840 acts to limit growth under prolonged iron limitation, a phenomenon likely adaptive for Mtb survival, similar to the slowed growth also observed with other stressors such as acidic pH and NO (11–15).

**Fig. 2.**
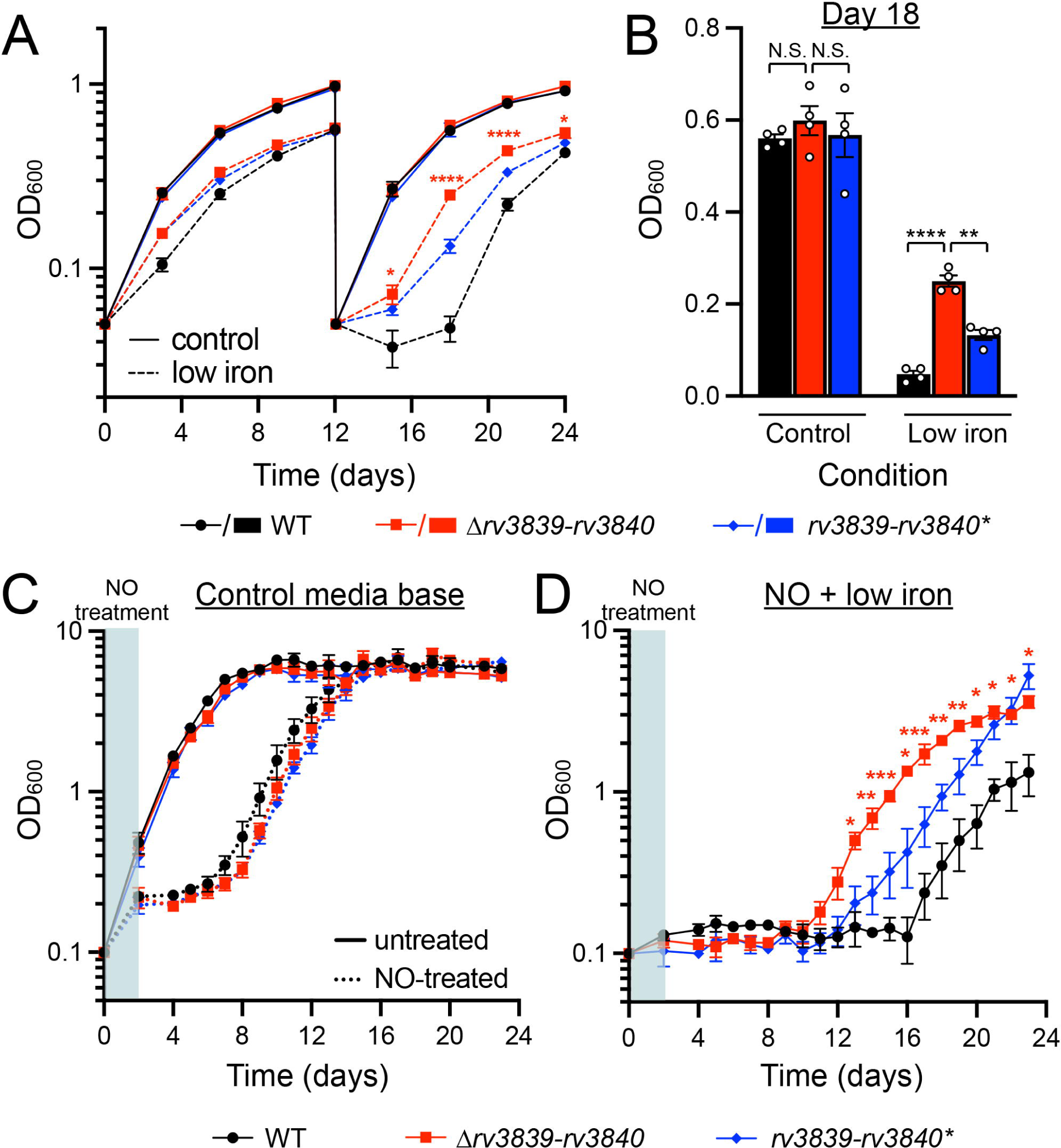
Rv3839-Rv3840 is important for maintenance of NO and iron stress-induced growth arrest. (A and B) Δ*rv3839-rv3840* Mtb fails to limit growth under iron limitation. Growth curves (A) and day 18 timepoint (B) of WT, Δ*rv3839-rv3840,* and *rv3839-rv3840\** (complemented strain) Mtb sequentially cultured in 7H9, pH 7.0 media (control) or iron-depleted minimal media with 100 µM 2’2’-dipyridyl (low iron). Bacterial growth was tracked by OD_600_ every 3 days. At day 12, the strains were sub-cultured into the same medium. Data are shown as means ± SEM from 4 experiments. p-values were obtained with unpaired t-tests with Welch’s correction in (A), comparing Δ*rv3839-rv3840* to WT Mtb in the low iron condition. p-values were obtained with a two-way ANOVA with Tukey’s multiple comparisons test in (B). N.S. not significant, * p<0.05, ** p<0.01, **** p<0.0001. (C and D) Δ*rv3839-rv3840* Mtb prematurely exits NO and low iron stress-induced growth arrest. WT, Δ*rv3839-rv3840,* and *rv3839-rv3840\** Mtb were grown in aerated conditions in 7H9, pH 7.0 and sub-cultured in either 7H9, pH 7.0 ± 6 doses of 100 µM DETA NONOate (C) over 30 hours (shaded area), or in iron-depleted minimal media with 100 µM 2’2’-dipyridyl and treated with 6 doses of 100 µM DETA NONOate (D) over 30 hours (shaded area). Bacterial growth was tracked by OD_600_ every day for 24 days. Data are shown as means ± SEM from 3 experiments. p-values in (D) were obtained with unpaired t-tests with Welch’s correction, comparing Δ*rv3839-rv3840* to WT Mtb. * p<0.05, ** p<0.01, *** p<0.001.

Given the augmented induction of *rv3839-rv3840* in the simultaneous presence of NO stress and iron limitation (Fig. 1), we next investigated whether deletion of *rv3839-*rv3840 also affected Mtb growth under the dual conditions of extended NO exposure and iron limitation. Mtb cultures were treated with 6 doses of 100 µM DETA NONOate, a NO donor with a half-life of ∼20 hours, over the course of 30 hours, to drive Mtb entry into a state of growth arrest (8). In the presence of extended NO exposure alone, Δ*rv3839-rv3840* Mtb exited growth arrest at the same time as WT Mtb (Fig. 2C). In contrast, Δ*rv3839-rv3840* Mtb strikingly exited growth arrest induced by simultaneous iron and extended NO stress significantly earlier than WT Mtb, with complementation partially restoring the WT outcome (Fig. 2D). This result indicates that Rv3839*-*Rv3840 impacts the ability of Mtb to maintain growth arrest under NO stress and iron limitation.

### Deletion of *rv3839-rv3840* alters Mtb cell length in the presence of NO and iron limitation

Remodeling of the Mtb cell envelope is thought to accompany the shift into growth arrest driven by NO and hypoxia (37–40). In addition, iron deprivation reduces cell wall thickness and increases susceptibility to cell membrane targeting antibiotics in *Mycobacterium smegmatis* (41, 42). Notably, *rv3840* encodes one of four LytR-CpsA-Psr (LCP) proteins in Mtb, others of which act in bacterial cell envelope maintenance by catalyzing the crosslinking reaction between arabinogalactan (AG) and peptidoglycan (PG) (43–47) and influence Mtb virulence and antibiotic resistance (43, 45–47). However, Rv3840 differs from the three characterized LCP proteins in Mtb (Rv3484, Rv3267, Rv0822c) in that it lacks the N-terminal transmembrane domain and C-terminal LytR_C domain found in canonical LCP proteins (44, 48) and has only the central catalytic domain. We therefore next sought to examine if exposure to iron limitation and NO stress and deletion of *rv3839*-*rv3840* affected Mtb morphology. A first observation was that WT Mtb were more elongated upon exposure to iron-limiting conditions as compared to 7H9, pH 7.0 control media (3.64 ± 0.33 µm versus 2.10 ± 0.04 µm, p<0.0001) (Fig. 3A and B). Interestingly, when NO stress was present together with iron limitation, WT Mtb lost this elongated phenotype (2.67 ± 0.09 µm versus 3.64 ± 0.33 µm, p<0.05) (Fig. 3A and B). Δ*rv3839-rv3840* Mtb was elongated compared to WT Mtb under both iron limitation (4.60 ± 0.07 µm versus 3.64 ± 0.33 µm, p<0.05) and NO stress + iron limitation (4.56 ± 0.17 µm versus 2.67 ± 0.09 µm, p<0.0001), with length restored to WT levels by complementation (Fig. 3A and B). These results are in accord with Mtb modulation of its growth and/or division in response to NO and iron limitation as environmental stresses. The continued presence of an elongated phenotype for Δ*rv3839-rv3840* Mtb in NO + iron limitation further supports that Rv3839*-*Rv3840 contributes to appropriate adaptation of Mtb growth and/or division when both environmental stressors are experienced concurrently.

**Fig. 3.**
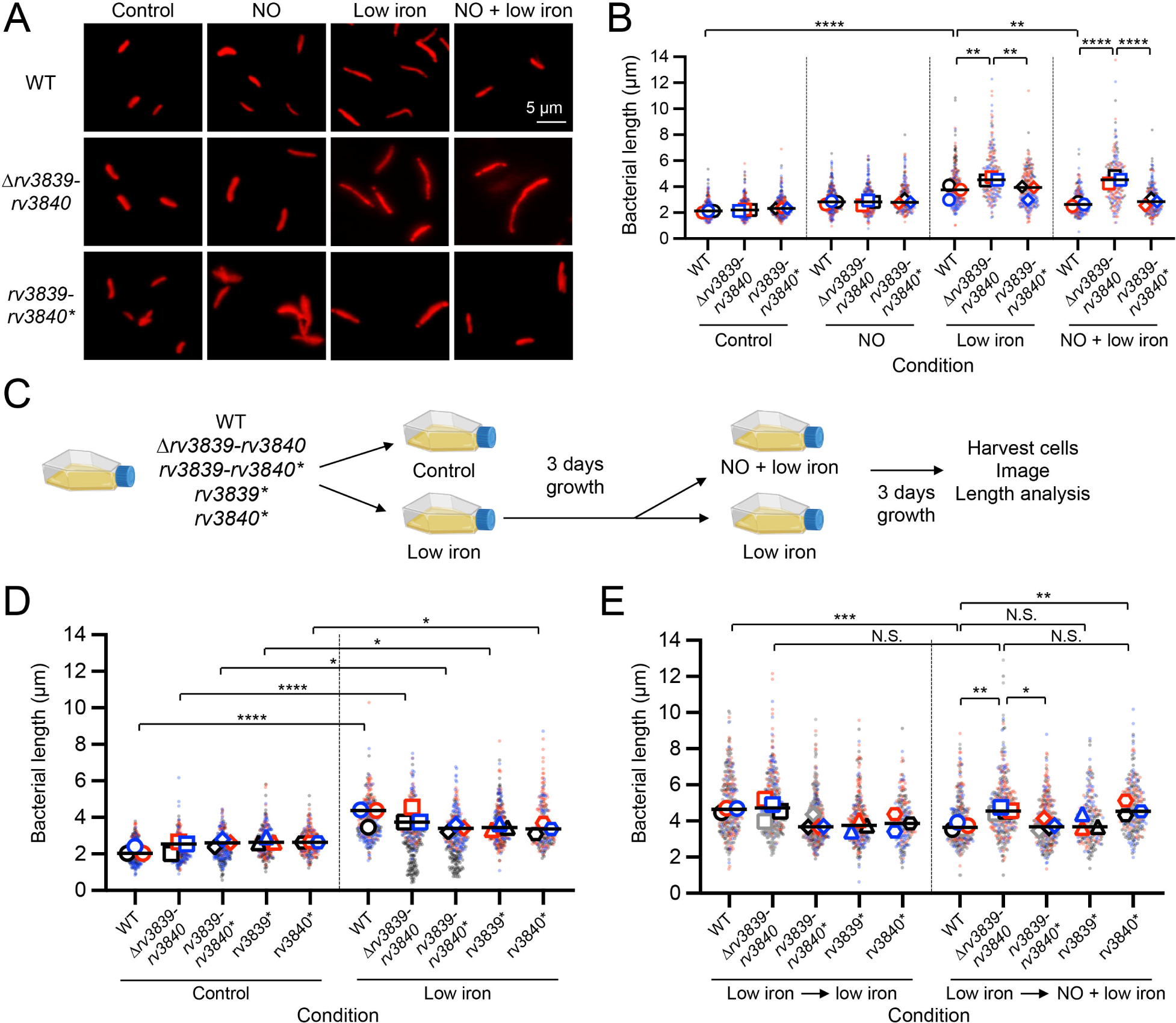
Deletion of *rv3839-rv3840* alters Mtb cell length in the presence of NO and iron limitation. (A and B) Δ*rv3839-rv3840* Mtb exhibit increased cell length in response to iron limitation and NO stress. WT, Δ*rv3839-rv3840*, and *rv3839-rv3840\** (complemented mutant) Mtb carrying a constitutive *smyc’*::mCherry reporter were sub-cultured into 7H9, pH 7.0 (control) or iron-depleted minimal media with 100 µM 2’2’-dipyridyl (low iron), ± 100 µM DETA NONOate for 3 days. (A) shows representative images, with lengths quantified in (B). Data are shown from 3 experiments. Each point represents a single bacterium and median values for each replicate are represented by the larger symbol. p-values were obtained with a two-way ANOVA with Tukey’s multiple comparisons test. ** p<0.01, **** p<0.0001. (C-E) Rv3839 regulates cell elongation under prolonged iron limitation and NO stress. (C) shows a schematic representation of the sequential low iron and NO exposure experiment. WT, Δ*rv3839-rv3840*, *rv3839-rv3840*, rv3839*,* and *rv3840\** Mtb carrying a constitutive *smyc’*::mCherry reporter were sub-cultured in 7H9, pH 7.0 (control) or iron-depleted minimal media with 100 µM 2’2’-dipyridyl (low iron). After 3 days, strains in the low iron media were sub-cultured into low iron medium ± 100 µM DETA NONOate for an additional 3 days. (D) shows bacterial lengths quantified after the initial 3 days of growth. (E) shows bacterial lengths quantified after the additional 3 days of growth in low iron media ± NO. Data are shown from 3-4 experiments. Each point represents a single bacterium and median values for each replicate are represented by the larger symbol. p-values were obtained with a two-way ANOVA with Tukey’s multiple comparisons test. N.S. not significant, * p<0.05, ** p<0.01, *** p<0.001, **** p<0.0001.

To begin to delineate the role of Rv3839 versus Rv3840, we restored only one gene at a time to Δ*rv3839-rv3840* Mtb by complementation (*rv3839\** or *rv3840\**). Additionally, here we introduced NO stress only after an initial 3 days of iron limitation (Fig. 3C), rather than simultaneously (as in Fig. 3B), to determine the impact of subsequent NO stress exposure on the elongated phenotype observed under iron limitation. WT, Δ*rv3839-rv3840, rv3839-rv3840*, rv3839*,* and *rv3840\** Mtb were grown in either 7H9, pH 7.0 medium (control), or in iron-limited medium to induce the elongated phenotype (Fig. 3D). Strains in the iron-limited condition were then sub-cultured into iron-limited medium ± 100 μM DETA NONOate (Fig. 3C). All strains remained elongated when sub-cultured from iron-limiting conditions into low iron medium again (Fig. 3E, left side of graph). However, subsequent introduction of NO stress decreased WT Mtb length (3.69 ± 0.09 µm, “low iron→ low iron + NO”, versus 4.61 ± 0.06 µm, “low iron→ low iron”, p<0.001) (Fig. 3E). In contrast, Δ*rv3839-rv3840* Mtb remained elongated in all conditions where iron limitation was present (4.66 ± 0.27 µm, “low iron→ low iron”), including upon introduction of NO stress (4.57 ± 0.07 µm, “low iron→ low iron + NO”) (Fig. 3E). These results suggest that Δ*rv3839-rv3840* Mtb is unable to appropriately respond to NO stress under iron-limiting conditions, in accord with the observed premature exit from growth arrest in NO + iron-limiting conditions (Fig. 2D). Surprisingly, complementation with *rv3839* alone restored cell length to WT levels under NO stress and iron limitation (3.90 ± 0.24 µm compared to 3.69 ± 0.09 µm), while complementation with *rv3840* did not (4.66 ± 0.24 µm compared to 3.69 ± 0.09 µm, p<0.01) (Fig. 3E). These data support that Rv3839 function affects Mtb morphology under NO stress and iron limitation, not Rv3840, as may have been expected *a priori* given its association with the LCP family of proteins.

### Endogenous heme biosynthesis contributes to maintenance of growth arrest in response to iron limitation

Rv3839 contains a domain of unknown function (DUF2470) found in heme-related proteins in bacteria and eukaryotes and is thought to be a heme-binding regulatory domain (49). For example, DUF2470 is found in HugZ, a heme oxygenase involved in iron release and uptake in *Helicobacter pylori* (36), and in GluBP, a regulatory protein that enhances heme synthesis in *Arabidopsis thaliana* by binding glutamyl-tRNA reductase (50, 51). Phylogenetic analysis has shown that genes encoding DUF2470-containing proteins are often found near genes related to iron homeostasis, suggesting a conserved role in iron and heme homeostasis (49). Given the primary role of Rv3839 observed above and that Rv3839 contains DUF2470, we thus next pursued further study of the role of the operon in endogenous heme biosynthesis. In Mtb, heme is synthesized via the coproporphyrin-dependent (CPD) pathway (52, 53). The initial steps of this pathway, 5-aminolevulinic acid (ALA) to coproporphyrinogen III, are shared with the protoporphyrin-dependent heme biosynthesis pathway used by most Gram-negative bacteria (Fig. 4A) (53, 54). To produce heme by the CPD pathway, coproporphyrinogen III is oxidized to coproporphyrin III, and then iron is inserted to form coproheme, which then undergoes double oxidative decarboxylation to yield heme (Fig. 4A) (53, 54). Given the requirement of iron in this process, we hypothesized that *rv3839-rv3840* is upregulated to dampen heme biosynthesis and prevent the production of heme when there is insufficient iron available.

**Fig. 4.**
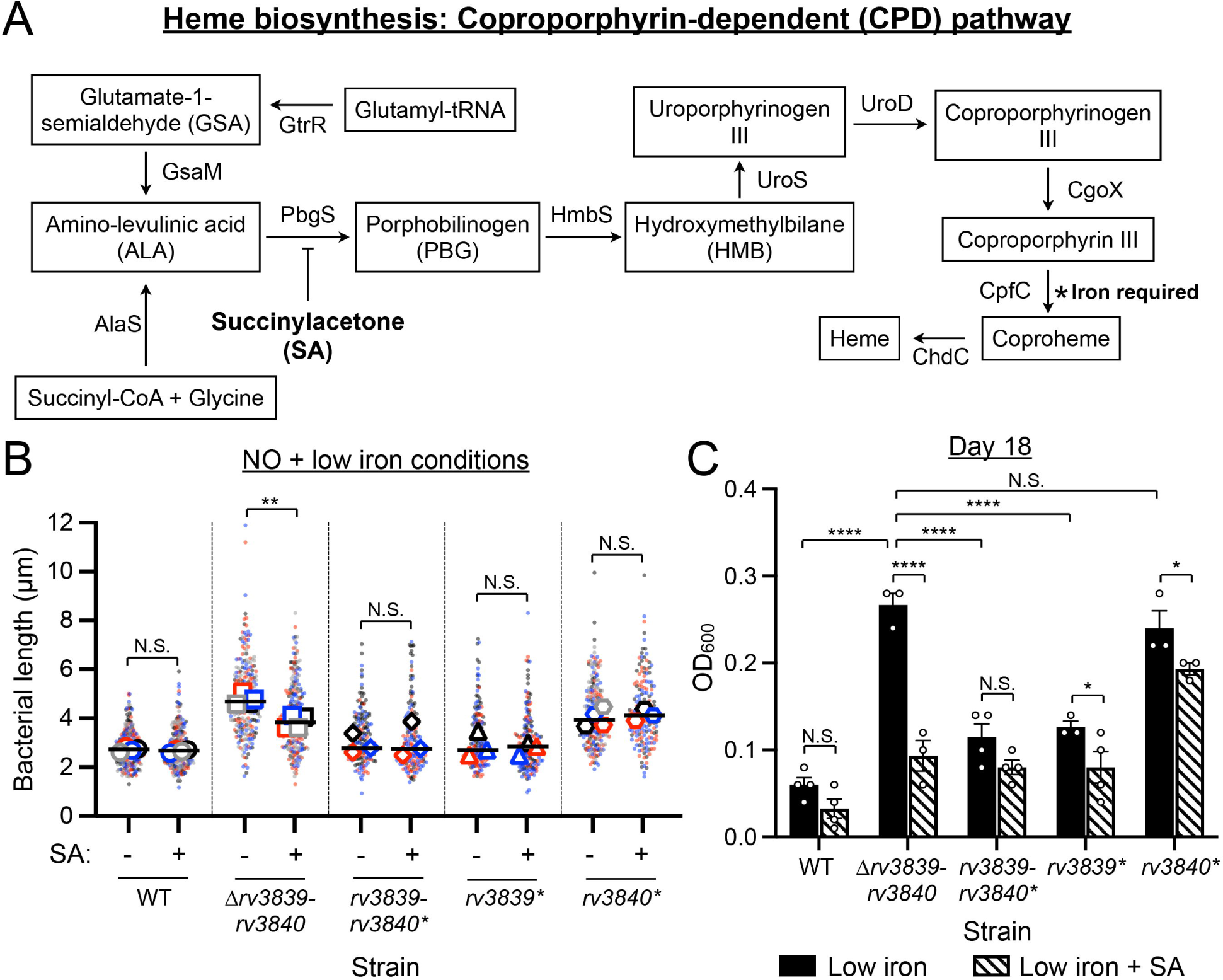
Inhibition of endogenous heme biosynthesis abrogates cell elongation and growth phenotypes of Δ*rv3839-rv3840* Mtb. (A) Schematic representation of Mtb endogenous heme biosynthesis pathway. (B) Succinylacetone (SA) treatment inhibits elongation of Δ*rv3839-rv3840* Mtb under NO + low iron conditions. WT, Δ*rv3839-rv3840*, *rv3839-rv3840*, rv3839*,* and *rv3840\** Mtb carrying a constitutive *smyc’*::mCherry reporter were sub-cultured in iron-depleted minimal media with 100 µM 2’2’-dipyridyl and 100 µM DETA NONOate (NO + low iron), ± 500 µM SA for 3 days. Fixed samples were analyzed via microscopy and bacterial cell length quantified. Data are shown from 3-4 experiments. Each point represents a single bacterium and median values for each replicate are represented by the larger symbol. p-values were obtained with a two-way ANOVA with Tukey’s multiple comparisons test. N.S. not significant, ** p<0.01. (C) Increased growth of Δ*rv3839-rv3840* Mtb under iron limitation is inhibited by SA treatment. WT, Δ*rv3839-rv3840*, *rv3839-rv3840*, rv3839*,* and *rv3840\** Mtb strains were cultured in iron-depleted minimal media with 100 µM 2’2’-dipyridyl ± 250 µM SA. At day 12, the strains were sub-cultured in the same medium. OD_600_ of the bacterial cultures at day 18 are shown. Data are shown as means ± SEM from 3-4 biological replicates. p-values were obtained with a two-way ANOVA with Tukey’s multiple comparisons test. N.S. not significant, * p<0.05, ** p<0.01, **** p<0.0001.

To study the impact of heme homeostasis on Mtb growth under NO stress and iron limitation, and the related role of Rv3839*-*Rv3840, we perturbed the endogenous heme biosynthesis pathway in Mtb with the inhibitor succinylacetone (SA). This compound blocks the activity of porphobilinogen synthase (PbgS) that converts ALA to porphobilinogen, one of the early steps in heme synthesis (Fig. 4A) (55, 56). We reasoned that if Rv3839-Rv3840 activates heme biosynthesis, WT Mtb would behave like Δ*rv3839-rv3840* Mtb if the pathway is blocked via SA treatment. If instead Rv3839-Rv3840 represses heme synthesis, blocking the pathway would restore Δ*rv3839-rv3840* Mtb cell length and growth to that of WT Mtb. We first investigated whether the elongated phenotype observed in Δ*rv3839-rv3840* Mtb under NO stress and iron limitation was dependent on increased flux through the heme biosynthesis pathway. Treatment of Δ*rv3839-rv3840* Mtb with SA resulted in a decrease in cell length under NO stress and iron limitation, compared to untreated samples (3.85 ± 0.14 µm compared to 4.75 ± 0.11 µm, p<0.001) (Fig. 4B). WT Mtb cell length was not impacted by SA treatment (Fig. 4B). These data support that endogenous heme biosynthesis impacts Mtb cell length and suggests that Rv3839-Rv3840 inhibits this pathway under NO stress and iron limitation.

We next investigated whether dysregulated heme biosynthesis in Δ*rv3839-rv3840* Mtb is a contributing factor to its inability to limit growth under iron limitation. Under iron-limiting conditions, Δ*rv3839-rv3840* Mtb exhibited increased growth compared to WT Mtb (Fig. 2A and 4C). Complementation with *rv3839*, but not *rv3840,* largely returned growth to WT levels (Fig. 4C). Blocking endogenous heme biosynthesis by treatment with SA under iron limitation resulted in growth inhibition of Δ*rv3839-rv3840* Mtb, restoring growth to levels more similar to WT in iron-limited medium (Fig. 4C). These data suggest that reduced heme biosynthesis contributes to Mtb growth arrest under iron limitation and that Rv3839 may act as an inhibitor of heme biosynthesis to maintain an adaptive growth-arrested state.

### Deletion of *rv3839-rv3840* results in accumulation of coproporphyrin III trimethyl ester in iron-limiting conditions

If Rv3839*-*Rv3840 indeed represses the heme biosynthesis pathway when iron is unavailable for coproheme production, we would expect a buildup of the upstream intermediates of the pathway in Δ*rv3839-rv3840* Mtb under iron limitation. We capitalized on the natural fluorescence of porphyrins to measure flux through the heme biosynthesis pathway in Δ*rv3839-rv3840* Mtb under iron limitation (57). Δ*rv3839-rv3840* Mtb exhibited strongly increased levels of free porphyrins when grown under iron limitation compared to WT Mtb (2696.67 ± 340.85 AFU versus 283.50 ± 14.91 AFU) (Fig. 5A). Porphyrin levels were dampened in Δ*rv3839-rv3840* Mtb under simultaneous NO stress and iron limitation, although levels were still elevated compared to WT Mtb (947.50 ± 45.05 AFU versus 287.17 ± 8.40 AFU) (Fig. 5A). Interestingly, either *rv3839* or *rv3840* was sufficient for complementation of the Δ*rv3839-rv3840* Mtb phenotype in this case (Fig. 5A), indicating that an increase in free porphyrins is not the sole driver of the elongated cell and growth phenotypes observed in Δ*rv3839-rv3840* Mtb. Together, these results support that under iron limitation, Rv3839-Rv3840 prevents the buildup of porphyrin intermediates.

**Fig. 5.**
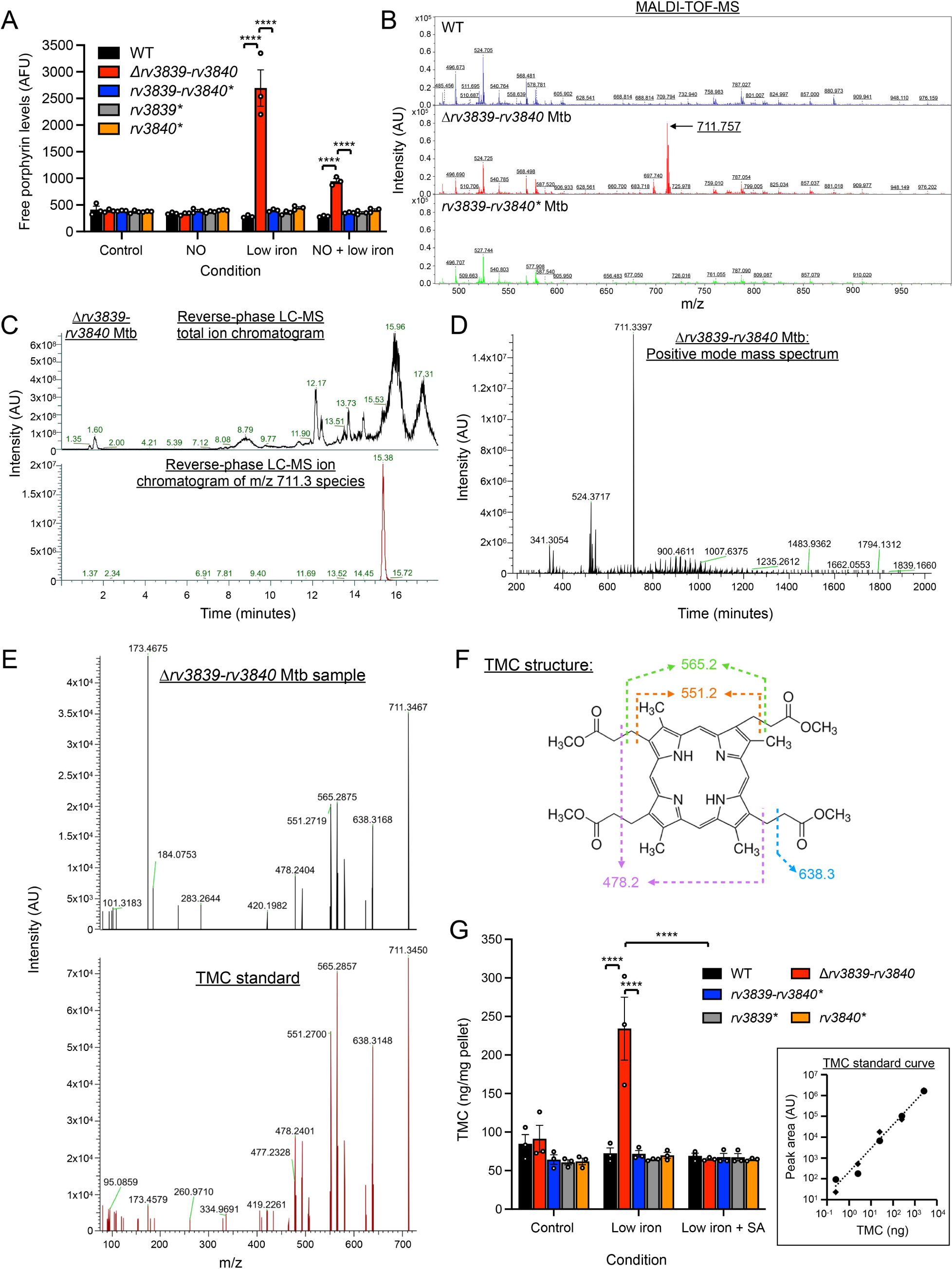
Deletion of *rv3839-rv3840* results in accumulation of coproporphyrin III tetramethyl ester (TMC) under iron limiting conditions. (A) Δ*rv3839-rv3840* Mtb has elevated levels of porphyrins under iron limitation. WT, Δ*rv3839-rv3840*, *rv3839-rv3840\**, *rv3839\**, and *rv3840\** Mtb were sub-cultured and grown for 3 days in 7H9, pH 7.0 or iron-depleted minimal media with 100 µM 2’2’-dipyridyl, ± 100 µM DETA NONOate. Samples were tested for intrabacterial porphyrin levels. Data are shown as means ± SEM from 3 experiments. AFU = arbitrary fluorescence unit. p-values were obtained with a two-way ANOVA with Tukey’s multiple comparisons test. **** p<0.0001. (B) Reflective positive MALDI-TOF spectra of WT, Δ*rv3839-rv3840*, and *rv3839-rv3840\** grown in iron-depleted minimal media with 100 µM 2’2’-dipyridyl (low iron) for 3 days. Accumulation of a m/z 711.757 ion in Δ*rv3839-rv3840* Mtb is indicated. AU = arbitrary units. (C) Reverse phase LC-MS ion chromatogram of Δ*rv3839-rv3840* Mtb grown in low iron. Total ion chromatogram is shown in the top panel. Ion chromatogram of species matching m/z 711.3 (± 100 ppm), showing a single elution peak at 15.38 min, is shown in the bottom panel. (D) Positive-mode mass spectrum of Δ*rv3839-rv3840* at retention time of 15.38 min, showing the accumulating peak of m/z 711.3. (E) Identification of the m/z 711.3 ion as coproporphyrin III tetramethyl ester (TMC). Higher-energy collisional dissociation spectrum of the precursor ion (m/z 711.3) from Δ*rv3839-rv3840* Mtb fragmented by higher collisional dissociation at 55 V (top) compared to the fragmentation pattern of the TMC standard (bottom). (F) Chemical structure of TMC with the masses of expected fragments. (G) TMC accumulation in Δ*rv3839-rv3840* Mtb under iron limitation can be blocked by succinylacetone (SA) treatment. WT, Δ*rv3839-rv3840*, *rv3839-rv3840\**, *rv3839\**, and *rv3840\** Mtb were grown 7H9, pH 7.0 (control), or iron-depleted minimal media with 100 µM 2’2’-dipyridyl (low iron) ± 500 µM SA for 3 days, and samples analyzed via MALDI-TOF-MS. TMC was quantified using a standard curve generated using TMC standards (inset standard curve graph). Data are shown as mean ± SEM from 3 experiments. p-values were obtained with a two-way ANOVA with Tukey’s multiple comparisons test. **** p<0.0001.

To elucidate the identity of the porphyrin-containing compound that accumulates in Δ*rv3839-rv3840* Mtb in iron-limiting conditions, WT, Δ*rv3839-rv3840*, *rv3839-rv3840*, rv3839\**, and *rv3840\** Mtb were grown in iron-limited medium and samples extracted for analysis by positive ion mode MALDI-TOF mass spectroscopy (MS). This analysis revealed a strong signal with an *m/z* value of 711.757 in Δ*rv3839-rv3840* Mtb (Fig. 5B). This m/z closely matched with 711.34, the monoisotopic mass of the protonated form of coproporphyrin III tetramethyl ester (TMC). To determine its identity, we analyzed lipid extracts from WT and Δ*rv3839-rv3840* Mtb using reverse phase LC-MS. The 711 peak eluted at 15.38 min (Fig. 5C and D), and the mass spectrum at 15.38 min indicated the 711 peak as a major peak from the Δ*rv3839-rv3840* Mtb extract (Fig. 5C and D). To further confirm the identity of this peak, we fragmented the parental ion from the Δ*rv3839-rv3840* Mtb sample (Fig. 5D); this analysis demonstrated that the patterns of the fragmentation were nearly identical to those of an authentic TMC (Fig. 5E and F). Having determined its identity, we used MALDI-TOF MS for high-throughput analysis. First, we determined the dose-response curve of TMC in MALDI-TOF MS, demonstrating a linear range between 0.25 – 2500 ng per spot (Fig. 5G, inset). Using this standard curve, we quantified TMC abundance in lipid extracts from WT, Δ*rv3839-rv3840* and the various complementation Mtb strains, which showed high accumulation of TMC in iron-limiting conditions in Δ*rv3839-rv3840* Mtb (234 ± 71 ng/mg pellet) compared to WT (72 ± 12 ng/mg pellet) (Fig. 5G). TMC levels were complemented by either *rv3839* or *rv3840*, and SA treatment lowered TMC levels in Δ*rv3839-rv3840* Mtb to WT levels (Fig. 5G), indicating that Rv3839-Rv3840 regulates the heme biosynthesis pathway upstream of porphobilinogen synthase.

### Rv3839-Rv3840 regulates Mtb heme homeostasis

Often, elevated intracellular porphyrin levels correspond with increased heme production. However, given the modification of coproporphyrin III to TMC, it was unclear whether TMC abundance was indicative of elevated heme production or whether this modification is a strategy for diverting excess precursors to prevent excess heme production under iron limiting conditions. To distinguish between these two possibilities, we measured total intracellular heme in WT, Δ*rv3839-rv3840*, *rv3839-rv3840*, rv3839\**, and *rv3840\** Mtb grown in iron-limiting conditions, with or without NO stress. WT Mtb showed a trend towards decreased heme levels under iron limitation compared to control media, and a further reduction in total heme with the addition of NO stress, although this did not reach statistical significance (Fig. 6A). In contrast, Δ*rv3839-rv3840* Mtb showed increased heme levels compared to WT Mtb under iron limitation (5146.83 ± 438.32 AFU versus 1088.00 ± 70.36 AFU, p<0.0001) and increased heme levels compared to WT Mtb under NO stress and iron limitation (3240.83 ± 349.66 versus 859.17 ± 78.05 AFU, p<0.0001) (Fig. 6A). Total heme levels were reduced in Δ*rv3839-rv3840* Mtb under the dual NO + iron limitation condition compared to iron limitation alone (Fig. 6A). Total heme levels were complemented by *rv3839* or *rv3839-rv3840*, but complementation with *rv3840* only partially reduced total heme levels compared to Δ*rv3839-rv3840* (Fig. 6A).

**Fig. 6.**
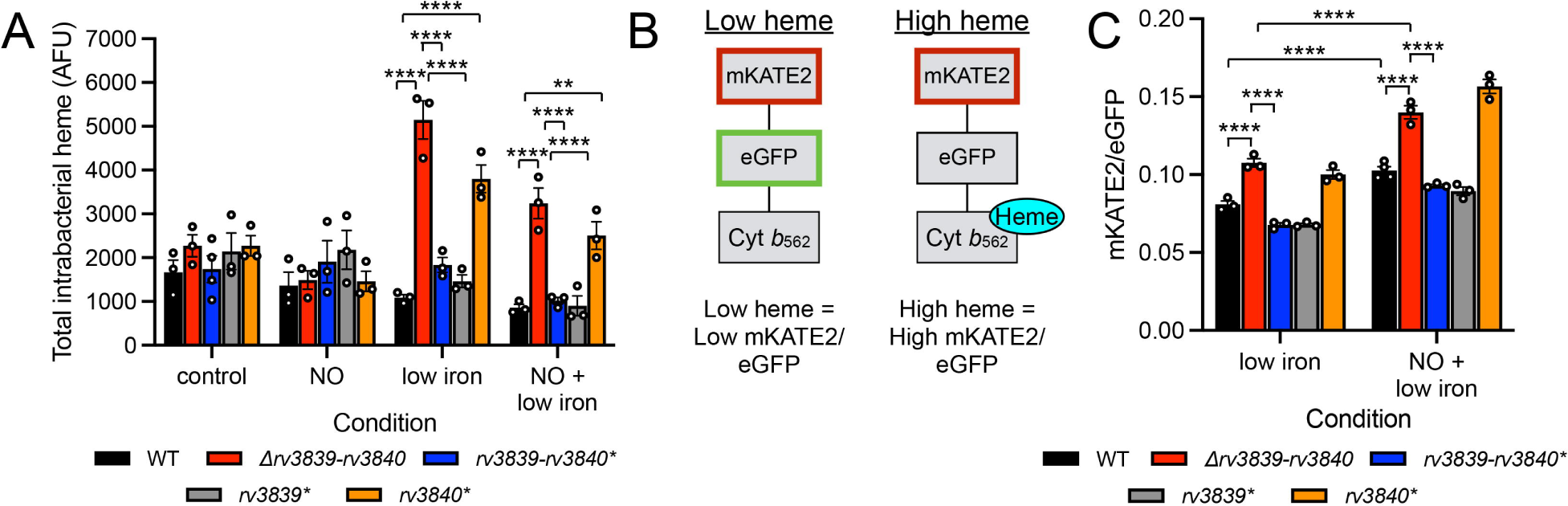
Rv3839-Rv3840 regulates intrabacterial heme levels. (A) Total cellular heme levels are elevated in Δ*rv3839-rv3840* Mtb under iron limitation ± NO stress. WT, Δ*rv3839-rv3840*, *rv3839-rv3840*, rv3839*,* and *rv3840\** Mtb were sub-cultured and grown for 3 days in 7H9, pH 7.0 (control) or iron-depleted minimal media with 100 µM 2’2’-dipyridyl (low iron) ± 100 µM DETA NONOate. Total cellular heme is calculated as fluorescence of treated samples with background porphyrin fluorescence subtracted. Data are shown as means ± SEM from 3 experiments. AFU = arbitrary fluorescence units. p-values were obtained with a two-way ANOVA with Tukey’s multiple comparisons test. ** p<0.01, **** p<0.0001. (B) Schematic representation of intrabacterial heme reporter design. Heme binding to cytochrome *b*_562_ (Cyt *b*_562_) results in quenching of eGFP fluorescence but does not impact mKATE2 fluorescence. (C) Intrabacterial labile heme levels are elevated in Δ*rv3839-rv3840* Mtb under iron limitation ± NO stress. WT, Δ*rv3839-rv3840*, *rv3839-rv3840*, rv3839*,* and *rv3840\** Mtb were sub-cultured and grown for 3 days in iron-depleted minimal media with 100 µM 2’2’-dipyridyl (low iron) ± 100 µM DETA NONOate. Intrabacterial labile heme is reported as the ratio of mKATE2/eGFP fluorescence, and data are shown as means ± SEM from 3 experiments. p-values were obtained with a two-way ANOVA with Tukey’s multiple comparisons test. N.S. not significant, **** p<0.0001.

Total intracellular heme is composed of both inert heme tightly bound to hemoproteins and a smaller fraction of kinetically labile heme that is more available for exchange and biological processes (57, 58). As an independent manner to assess intrabacterial heme levels, we introduced a labile heme reporter into Mtb, where fluorescence of green fluorescent protein (GFP) is quenched upon binding of labile heme to cytochrome *b*_562_, with a constitutively expressed mKATE2 enabling ratiometric analysis (Fig. 6B) (57, 59). As with total intrabacterial heme, Δ*rv3839-rv3840* Mtb showed increased intrabacterial labile heme levels compared to WT Mtb under both iron limitation (0.108 ± 0.003 versus 0.081 ± 0.002 mKATE2/eGFP signal, p<0.0001) and NO stress and iron limitation combined (0.140 ± 0.004 versus 0.103 ± 0.003 mKATE2/eGFP signal, p<0.0001) (Fig. 6C). The increase in labile heme in Δ*rv3839-rv3840* Mtb was lower than the increase in total intrabacterial heme observed, likely due to labile heme representing a lower overall proportion of the heme present in the bacteria. Interestingly, WT Mtb showed elevated labile heme levels under NO stress and iron limitation compared to iron limitation alone (0.103 ± 0.003 versus 0.081 ± 0.002 mKATE2/eGFP signal, p<0.0001) (Fig. 6C), suggesting that NO stress induces increased *de novo* synthesis of heme, even as total intrabacterial heme levels decrease (Fig. 6A). Labile heme levels were complemented by *rv3839* or *rv3839-rv3840*, but complementation with *rv3840* did not reduce labile heme levels compared to Δ*rv3839-rv3840* (Fig. 6C). Together, these data support that the abundance of TMC in Δ*rv3839-rv3840* Mtb under iron limitation corresponds with increased total intracellular heme, and demonstrate a role for Rv3839-Rv3840 as a repressor of heme synthesis that is important in Mtb adaptation to NO stress and iron limitation.

## DISCUSSION

The ability of Mtb to sense and integrate its response to multiple environmental cues is critical for its survival in the host environment. Here, we show that the response of Mtb to NO stress is augmented in the simultaneous presence of iron limitation and vice versa, and reveal an important role of Rv3839*-*Rv3840 in the regulation of heme biosynthesis and the adaptive response of the bacterium to iron limitation and NO stress. We propose a model wherein the *rv3839-rv3840* operon is upregulated under NO stress and iron limitation to repress heme biosynthesis in Mtb due to the reduced availability of iron for heme production (Fig. 7). Loss of this regulation upon deletion of *rv3839-rv3840* leads to the accumulation of modified porphyrin intermediates and intracellular heme, resulting in an elongated cell phenotype and premature exit from NO and iron limitation-induced adaptive growth arrest (Fig. 7).

**Fig. 7.**
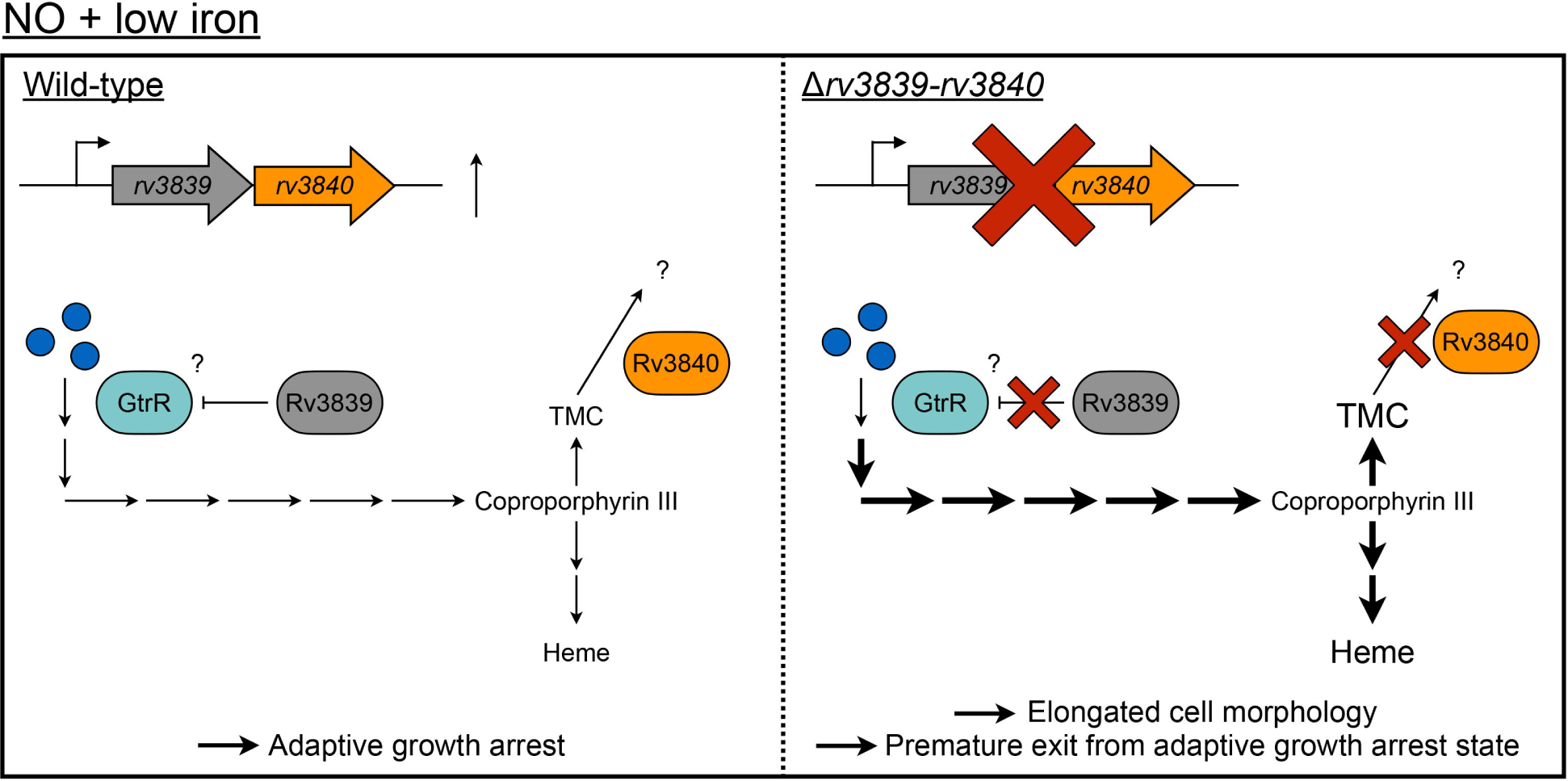
Model of Rv3839/Rv3840 regulation of heme biosynthesis. Under NO stress and low iron, WT Mtb (left side of the model) regulates heme biosynthesis to balance the requirement for heme in the NO response with the limitation imposed by the lack of iron. *rv3839/rv3840* are expressed and regulate heme biosynthesis early in the pathway, upstream of porphobilinogen synthase, and by modulating buildup of porphyrin intermediates respectively. Blue circles represent glutamyl-tRNA and other precursor molecules. This regulation of heme biosynthesis allows Mtb to maintain adaptive growth arrest. Deletion of *rv3839-rv3840* (right side of the model) results in increased flux through the heme biosynthetic pathways, resulting in the buildup of modified porphyrin intermediates and intracellular heme. The dysregulation of the pathway results in elongated cell morphology and premature exit from an adaptive growth arrest state.

Our observation that Mtb exhibits elongated cell morphology under iron-limiting conditions is in accord with previous work showing that iron starvation inhibits late-stage cytokinesis in *Escherichia coli* (60). We thus posit that the elongated cell morphology observed in Mtb upon iron limitation reflects a similar inhibition of cytokinesis; introduction of NO stress halts growth entirely and results in a shorter cell length, while deletion of *rv3839-rv3840* prevents appropriate sensing of NO stress in the context of iron limitation. Interestingly, our results show that it is Rv3839, not Rv3840, that is a driver of the elongated cell phenotype. As noted above, Rv3840 is an atypical LCP protein as it lacks the N-terminal transmembrane domain and C-terminal LytR_C domain found in canonical LCP protein domain structure (46, 48). In addition to the three canonical LCP proteins, Mtb encodes two LytR_C domain-only proteins, VirR (Rv0431) and Cei (Rv2700). Cei contributes to Mtb cell envelope integrity and virulence (61), and loss of VirR function results in increased vesiculogenesis (62, 63), a phenotype also associated with iron limitation (64). The existence of these two LytR_C domain-only proteins alongside Rv3840 raises the intriguing possibility that these proteins interact to reconstitute a “complete” LCP protein with function at the Mtb cell envelope. Additionally, LCP proteins are known to exhibit functional redundancy in Mtb (43) and other organisms (43, 65, 66), and have been shown to be essential for septum formation and cell separation in *Staphylococcus aureus* and *Streptococcus pneumoniae* (66, 67). Future studies analyzing possible interplay between Rv3840 and VirR and/or Cei, as well as the impact of NO stress and iron limitation on bacterial growth and cell division, will shed light on the involvement of LCP proteins in these processes.

One of the striking phenotypes associated with deletion of *rv3839-rv3840* is its premature exit from NO and iron limitation-induced growth arrest. Although TMC has been shown to accumulate in dormant *M. smegmatis* under acidified conditions (68, 69), *M. smegmatis* is a fast-growing mycobacterial species and has not been shown to enter into NO-induced growth arrest (70). It is also unclear why coproporphyrin III is methylated in Δ*rv3839-rv3840* Mtb and what purpose this modification serves. Given that WT Mtb does not normally accumulate TMC under NO and iron limitation stress, it seems unlikely that modification of coproporphyrin III and its accumulation would be protective. Indeed, SA treatment of the *rv3840\** complement Mtb strain under iron limitation does not dampen growth to the same degree as Δ*rv3839-rv3840* Mtb, although the treatment does reduce TMC levels back to those of WT Mtb. Notably, our results show that Δ*rv3839-rv3840* Mtb is unable to properly respond to the introduction of NO stress in the context of iron limitation as evidenced by its elongated cell morphology. Mtb has multiple heme enzymes with important roles, including the NO/hypoxia sensors DosS and DosT (31), truncated hemoglobin (TrHbN) (32–34), catalase peroxidase (KatG) (71), and cytochrome P450s (CYPs) (72), all of which may be impacted by the dysregulation of intrabacterial heme synthesis in Δ*rv3839-rv3840* Mtb, resulting in premature exit from an adaptive state of growth arrest. Mtb also encodes a heme-containing bacterioferritin, BfrA, which is required for efficient storage of iron under iron limiting conditions, and previous work has demonstrated that the heme-bound form of BfrA exhibits enhanced iron release compared to heme-free variants (73, 74). Increased intrabacterial heme levels in Δ*rv3839-rv3840* Mtb may therefore cause increased release of iron from storage proteins, resulting in enhanced growth under NO stress and iron limitation.

Another outstanding question from this work is the separate functional roles of Rv3839 versus Rv3840 and the precise mechanism by which they regulate heme biosynthesis. In *Corynebacterium glutamicum,* it has been shown that transcription of heme synthesis proteins is controlled in part by the iron-dependent regulator, DtxR (52, 75, 76). However, in mycobacteria, regulation of heme biosynthesis by iron or other factors has been more ambiguous. Glutamate semialdehyde aminomutase (GsaM) and ferrochelatase (CpfC), two enzymes in the heme biosynthesis pathway, were shown to have decreased expression in iron limiting media (26). In contrast, porphobilinogen synthase (PbgS) was upregulated under those same conditions (26). In our work, we have seen no evidence that deletion of *rv3839-rv3840* impacts the transcript levels of heme biosynthesis proteins, suggesting that Rv3839*-*Rv3840 may instead impact protein levels or activity state. Our results show that complementation with *rv3839* alone is sufficient to restore Δ*rv3839-rv3840* Mtb cell length, intrabacterial heme levels, and growth to that of WT Mtb, supporting Rv3839 as the main driver of the NO stress and iron-limitation-related phenotypes. Further, regulation of the heme biosynthesis pathway by Rv3839 likely occurs upstream of PbgS, as inhibition with SA dampens the growth of Δ*rv3839-rv3840* under iron limitation. We posit that Rv3839 regulates the activity of glutamyl-tRNA reductase (GtrR), which serves as a regulatory node for the heme biosynthesis pathway in multiple organisms (52, 75–78) and has been shown to be regulated by other DUF2470-containing proteins (50, 51). Our results also support a functional role for Rv3840 in preventing the buildup of precursor molecules in the heme biosynthesis pathway, as complementation with *rv3840* is sufficient to restore TMC but not heme levels to that of WT Mtb. These results support a model in which Rv3839 functions to repress heme biosynthesis early in the pathway, while Rv3840 acts as a secondary layer of regulation downstream to prevent the buildup of porphyrin intermediates (Fig. 7). Further experiments will be required to delineate the molecular mechanisms underlying how Rv3839 and Rv3840 regulate heme biosynthesis.

NO stress and iron limitation are two major environmental stressors Mtb encounters during infection, and the bacterium mounts a robust transcriptional response in order to adapt and survive. Our findings here emphasize the integrated nature of the NO and iron limitation stress responses and identify Rv3839 and Rv3840 as key factors involved in the regulation of heme biosynthesis in response to these two signals. Mtb response to NO stress requires iron-containing prosthetic groups, including heme, for proper sensing and response, even while NO simultaneously degrades factors such as Fe-S clusters. Therefore, the allocation of these resources is likely tightly regulated by the bacteria, with Rv3839*-*Rv3840 acting as one such layer of regulation. Our work adds to the understanding of how heme biosynthesis is regulated, and we propose that further studies investigating how the integration of NO stress, iron limitation, and heme biosynthesis impact Mtb cell division and growth will continue to reveal important insight into how Mtb environmental response is coordinated with its replication control.

## MATERIALS AND METHODS

### Mtb strains and culture

Mtb CDC1551 was used as the parental strains for all assays, and Mtb strains were cultured and maintained as previously described, with 7H9 broth supplemented with 10% OADC, 0.2% glycerol, 0.05% Tween 80, and 100 mM MOPS used for buffering to pH 7.0 (12). Iron-depleted minimal media was made as previously described (8, 26). All antibiotics were added as appropriate at the following concentrations: 100 μg/ml streptomycin, 50 μg/ml hygromycin, 50 μg/ml apramycin, and 25 μg/ml kanamycin. Δ*rv3839-rv3840* Mtb and its complements were constructed with methods as previously described (4), with the Δ*rv3839-rv3840* mutation consisting of a deletion beginning at nucleotide 107 of the *rv3839* open reading frame through the *rv3840* stop codon. The *rv3839-rv3840* complement (*rv3839-rv3840\**) consisted of a region beginning 830 bp upstream of the *rv3839* start codon and included both the *rv3839* and *rv3840* open reading frames. The same promoter region was used to drive expression of *rv3839* or *rv3840* in the single gene complements (*rv3839\** and *rv3840\**). The *smyc’*::mCherry (4) and intrabacterial heme (HS1) reporters (57, 59, 79) introduced to indicated strains were previously described.

### RNA sequencing and qRT-PCR analysis

For RNA sequencing (RNAseq) and qRT-PCR analyses, log-phase Mtb cultures (OD_600_ ∼0.6) grown in aerated conditions were used to inoculate filter-capped T75 flasks laid flat at an OD_600_ = 0.3, containing 12 ml: (i) iron-depleted minimal medium + 150 µM FeNO_3_ (control condition), (ii) iron-depleted minimal medium + 150 µM FeNO_3_ + 100 µM DETA NONOate (NO condition), (iii) iron-depleted minimal medium + 100 µM 2’2’-dipyridyl (low iron condition), or (iv) iron-depleted minimal medium + 100 µM 2’2’-dipyridyl + 100 µM DETA NONOate (NO + low iron condition). A wash step with iron-depleted minimal medium was performed prior to inoculation of the culture into the various test conditions. After four hours of exposure, Mtb samples were collected and RNA extracted as previously described (80). For RNAseq, three biological replicates were prepared for each condition, and library preparation was performed by the Tufts University Genomics Core Facility using the Illumina stranded total RNA with Ribo-Zero Plus kit. Barcoded samples were pooled and sequenced on an Illumina NovaSeq X Plus (single-end 100 bp reads). RNAseq data were analyzed as previously described (81). A log_2_-fold change ≥1 was used as the cutoff for the list of genes differentially expressed in various conditions in WT Mtb. A cutoff of log_2_-fold change ≥0.6 was used when comparing the effect of combined NO + low iron exposure versus either single condition (i.e. NO or low iron). qRT-PCR experiments were carried out and analyzed as previously described (82).

### Mtb growth assays

For low iron growth assays, log-phase Mtb strains were used to inoculate 10 ml of 7H9, pH 7.0 or iron-depleted minimal medium + 100 µM 2’2’-dipyridyl at a starting OD_600_ = 0.05 in standing, filter-capped T25 flasks. A wash step was conducted prior to inoculation as described above, and OD_600_ was measured at the indicated timepoints. After 12 days of growth, the strains were sub-cultured to an OD_600_ = 0.05 in the same media type. For growth assays testing the impact of succinylacetone (4,6-dioxoheptanoic acid, Sigma) treatment, strains were treated with 250 µM succinylacetone at days 0 and 12. NO and low iron growth arrest assays were conducted as previously described (8). In brief, log-phase Mtb cultures grown in aerated conditions were sub-cultured to OD_600_ = 0.1 in 12 ml 7H9, pH 7.0 or iron-depleted minimal medium + 100 µM 2’2’-dipyridyl, with a wash step as described above, in filter-capped T75 flasks laid flat. Cultures were then treated with 100 µM DETA NONOate 6 times across 30 hours, and growth was tracked over time via OD_600_ measurement.

### Mtb length measurements

To examine Mtb cell lengths, each of the strains carrying a constitutively expressed *smyc’*::mCherry reporter were grown under aerated conditions before inoculation at an OD_600_ = 0.1 in filter-capped T25 flasks laid flat containing 4 ml of: (i) 7H9, pH 7.0 (control condition), (ii) 7H9, pH 7.0 + 100 µM DETA NONOate (NO condition), (iii) iron-depleted minimal medium + 100 µM 2’2’-dipyridyl (low iron condition), or (iv) iron-depleted minimal medium + 100 µM 2’2’-dipyridyl + 100 µM DETA NONOate (NO + low iron condition). A wash step was carried out as described above prior to inoculation. Strains were grown for 3 days before samples were fixed in 4% paraformaldehyde (PFA) overnight and resuspended in phosphate-buffered saline (PBS) + 0.1% Tween 80.

For sequential exposure experiments, Mtb strains each carrying a constitutively expressed *smyc’*::mCherry reporter were grown under aerated conditions before inoculation, with a wash step, at an OD_600_ = 0.1 in filter-capped T25 flasks laid flat containing 4 ml of 7H9, pH 7.0 or iron-depleted minimal medium + 100 µM 2’2’-dipyridyl. After 3 days growth, the strains were sub-cultured to an OD_600_ = 0.1 in the same media type ± 100 µM DETA NONOate. After an additional 3 days growth, samples were fixed in 4% PFA overnight and resuspended in PBS + 0.1% Tween 80.

To test the impact of succinylacetone treatment on bacterial cell length, strains carrying a constitutively expressed *smyc’*::mCherry reporter were grown under aerated conditions before inoculation, with a wash step, at an OD_600_ = 0.1 in filter-capped T25 flasks laid flat containing 4 ml of iron-depleted minimal medium + 100 µM 2’2’-dipyridyl + 100 µM DETA NONOate (NO + low iron) ± 500 µM succinylacetone. Strains were grown for 3 days before samples were fixed in 4% PFA overnight and resuspended in PBS + 0.1% Tween 80.

For all length measurements, samples were mounted using ProLong Glass antifade (Invitrogen) and imaged as previously described (83). Bacterial lengths were measured in Volocity, with ≥50 cells measured per strain per condition for each biological replicate.

### Mass spectrometric analysis of lipid extracts

Log-phase Mtb strains grown in aerated conditions were sub-cultured to OD_600_ = 0.1 in 12 ml of: (i) 7H9, pH 7.0, (ii) 7H9, pH 7.0 + 100 µM DETA NONOate (NO condition), (iii) iron-depleted minimal medium + 100 µM 2’2’-dipyridyl (low iron condition), or (iv) iron-depleted minimal medium + 100 µM 2’2’-dipyridyl + 100 µM DETA NONOate (NO + low iron condition). Strains were grown for 3 days before pelleting and extraction with 100% methanol. Lipids were subsequently extracted from Mtb cell pellets by chloroform/methanol and purified by 1-butanol/water partitioning as previously described (84). Final butanol phase was dried on a speed-vac concentrator and resuspended at 1 mg wet cell pellet equivalent per µl of water-saturated 1-butanol.

For matrix-assisted laser desorption/ionization time-of-flight mass spectrometry (MALDI-TOF-MS), we first spotted 0.5 µl of matrix (a saturated solution of α-cyano-4-hydroxycinnamic acid prepared in 70% acetonitrile with 0.1% trifluoroacetic acid in water) on a polished steel MTP target plate (Bruker #8280781) and allowed it to air-dry. We then spotted 0.5 µl of the lipid extract on top of the dried matrix and allowed it to air-dry. Another 0.5 µl of the matrix solution was then placed on top of the lipid sample and allowed to air-dry. Mass spectra were acquired using a UltrafleXtreme MALDI-TOF/TOF mass spectrometer (Bruker Daltonics) operated in reflective positive-ion mode with the laser intensity set at 80%. Spectra were collected by averaging 2000 laser shots per sample. The standard coproporphyrin III tetramethyl ester (Sigma-Aldrich, C7157) was used to identify and quantify the mass peak of tetramethyl coproporphyrin.

For LC-MS, lipid extract (3 µl in water-saturated 1-butanol, purified from 1.5 mg wet cell pellet) was dried by a speed-vac concentrator and resuspended in 10 µl of 10% (v/v) acetonitrile in water (solvent A). The lipid suspension was transferred to an autosampler vial. HPLC separation and MS were performed using an Orbitrap Fusion Tribrid Mass Spectrometer (Thermo) with a Waters Acquity BEH C18 reversed-phase column (1.7 µm x 2.1 mm x 50 mm). Three µl was injected, and the column was eluted at 0.100 ml/min with a binary gradient from 0% to 100% solvent B (100% acetonitrile): 2–12 min (0 ➔100% B); 12–14 min (100 ➔ 0% B); 14–18 min (0% B). Spectra were collected in positive ion mode from *m/z* 197 to 2,000 at 1 spectrum/s. Internal calibration was performed with the EASY-IC system integral to the instrument. Fragmentation was performed on the peak of interest using higher energy collisional dissociation (HCD) at energies between 45-65V. Fragmentation data in Thermo Fisher Compound Discoverer were used to determine the molecular identity.

### Intrabacterial porphyrin and total heme measurements

Intrabacterial porphyrin and total heme measurements were conducted as previously described (57), with some modifications. Briefly, log-phase Mtb strains grown in aerated conditions were sub-cultured to OD_600_ = 0.1 in 12 ml of: (i) 7H9, pH 7.0 (control condition), (ii) 7H9, pH 7.0 + 100 µM DETA NONOate (NO condition), (iii) iron-depleted minimal medium + 100 µM 2’2’-dipyridyl (low iron condition), or (iv) iron-depleted minimal medium + 100 µM 2’2’-dipyridyl + 100 µM DETA NONOate (NO + low iron condition). After 3 days growth, samples were normalized to an OD_600_ = 2 and washed twice, first with UltraPure H_2_O (Invitrogen) + 0.1% Tween 80, then with UltraPure H_2_O, before resuspension in 1 ml PBS. After resuspension, 500 µl of cells were pelleted and frozen at -80°C overnight. For the assay, cell pellets were thawed and resuspended in 500 µl of 20 mM oxalic acid and stored at 4°C overnight. Then, 500 µl of 2 M oxalic acid was added to each cell suspension, and the sample divided into two 500 µl aliquots. One was stored in the dark at room temperature as a blank and to obtain the intrabacterial porphyrin measurements. The other set was boiled at 100°C for 30 minutes covered (total heme measurements). Both sets were centrifuged at 21,100 x *g* for 2 minutes to remove cell debris. For each sample, 200 µl of supernatant was plated in technical duplicate in a 96-well black flat-bottom plate, and fluorescence measured using a Biotek Synergy Neo2 multi-mode microplate reader with excitation at 400 nm and emission at 608 nm. Heme levels were calculated by subtracting fluorescence of intrabacterial porphyrin (blank samples) from fluorescence of boiled samples. Values are reported as arbitrary fluorescence units (AFUs).

### Intrabacterial heme reporter assay

Labile intrabacterial heme was measured using Mtb strains carrying the HS1 reporter as described previously (57, 59, 79). Briefly, strains were grown as in the intrabacterial porphyrin and total heme assay. After 3 days growth, samples were normalized to an OD_600_ = 1 in 500 µl PBS. To measure fluorescence, 200 µl of supernatant was plated in technical duplicate in 96-well black flat-bottom plate, and fluorescence was measured using a Biotek Synergy Neo2 multi-mode microplate reader. Excitation was 480 nm and emission at 510 nm for eGFP, with excitation at 580 nm and emission at 620 nm for mKATE2. Three reads were taken over 10 min to account for variability in fluorescence over time and averaged as one ratio.

### Accession number

RNA sequencing data has been deposited in the NCBI GEO database (ID number pending and will be added prior to publication).

## ACKNOWLEDGEMENTS

We thank Amit Reddi (Georgia Institute of Technology) for generously providing the HS1 intrabacterial heme reporter plasmid, and acknowledge support from the IALS Mass Spectrometry Core (Director, Stephen Eyles). We thank Yue Chen for help with RNAseq analysis, and all members of the Tan laboratory for helpful discussion. This work was supported by grants from the National Institutes of Health (NIH) (R01 AI143768 and R21 AI168597) to ST, and by grants from the Mizutani Foundation for Glycoscience and University of Massachusetts Amherst Institute for Applied Life Sciences (IALS) Core Facilities Incentive Fund to YSM. NFQ was supported in part by training grant T32 GM007310 from the National Institutes of Health. KNG was supported by a fellowship from the University of Massachusetts Amherst as part of the Chemistry-Biology Interface Training Program (National Research Service Award T32 GM139789). The funders had no role in study design, data collection and analysis, or preparation of the manuscript.

This manuscript is the result of funding in part by the NIH. It is subject to the NIH Public Access Policy. Through acceptance of this federal funding, NIH has been given a right to make this manuscript publicly available in PubMed Central upon the Official Date of Publication, as defined by NIH.

NFQ and ST conceptualized the project. NFQ, KNG, and YSM carried out experiments. ST and YSM supervised the work. ST and YSM were responsible for funding acquisition. NFQ and ST wrote the original draft of the manuscript, with final review and editing performed by all authors.

## SUPPLEMENTAL MATERIAL FIGURE LEGENDS

Table S1. Comparison of effect of iron limitation on genes differentially expressed ≥1 log_2_-fold in NO conditions in WT Mtb (4 hour exposure).

Table S2. Comparison of effect of NO on genes differentially expressed ≥1 log_2_-fold in low iron conditions in WT Mtb (4 hour exposure).

Table S3. Genes differentially expressed ≥1 log_2_-fold in NO + low iron conditions compared to control conditions in WT Mtb (4 hour exposure)

## Notes

### Competing Interest Statement

The authors have declared no competing interest.

### Summary of Updates

Supplemental files have been added in this revision.

## REFERENCES

1. Pai M, Behr MA, Dowdy D, Dheda K, Divangahi M, Boehme CC, Ginsberg A, Swaminathan S, Spigelman M, Getahun H, Menzies D, Raviglione M. 2016. Tuberculosis. Nat Rev Dis Primers 2:16076.

2. Cadena AM, Fortune SM, Flynn JL. 2017. Heterogeneity in tuberculosis. Nat Rev Immunol 17:691–702.

3. Lavin RC, Tan S. 2022. Spatial relationships of intra-lesion heterogeneity in *Mycobacterium tuberculosis* microenvironment, replication status, and drug efficacy. PLoS Pathog 18:e1010459.

4. Tan S, Sukumar N, Abramovitch RB, Parish T, Russell DG. 2013. *Mycobacterium tuberculosis* responds to chloride and pH as synergistic cues to the immune status of its host cell. PLoS Pathog 9:e1003282.

5. Gautam US, McGillivray A, Mehra S, Didier PJ, Midkiff CC, Kissee RS, Golden NA, Alvarez X, Niu T, Rengarajan J, Sherman DR, Kaushal D. 2015. DosS Is required for the complete virulence of *Mycobacterium tuberculosis* in mice with classical granulomatous lesions. Am J Respir Cell Mol Biol 52:708–16.

6. Park HD, Guinn KM, Harrell MI, Liao R, Voskuil MI, Tompa M, Schoolnik GK, Sherman DR. 2003. Rv3133c/dosR is a transcription factor that mediates the hypoxic response of *Mycobacterium tuberculosis*. Mol Microbiol 48:833–43.

7. Walters SB, Dubnau E, Kolesnikova I, Laval F, Daffe M, Smith I. 2006. The *Mycobacterium tuberculosis* PhoPR two-component system regulates genes essential for virulence and complex lipid biosynthesis. Mol Microbiol 60:312–30.

8. Giacalone D, Yap RE, Ecker AMV, Tan S. 2022. PrrA modulates *Mycobacterium tuberculosis* response to multiple environmental cues and is critically regulated by serine/threonine protein kinases. PLoS Genet 18:e1010331.

9. Haydel SE, Malhotra V, Cornelison GL, Clark-Curtiss JE. 2012. The *prrAB* two-component system is essential for *Mycobacterium tuberculosis* viability and is induced under nitrogen-limiting conditions. J Bacteriol 194:354–61.

10. Ohno H, Zhu G, Mohan VP, Chu D, Kohno S, Jacobs WR, Jr., Chan J. 2003. The effects of reactive nitrogen intermediates on gene expression in *Mycobacterium tuberculosis*. Cell Microbiol 5:637–48.

11. Voskuil MI, Schnappinger D, Visconti KC, Harrell MI, Dolganov GM, Sherman DR, Schoolnik GK. 2003. Inhibition of respiration by nitric oxide induces a *Mycobacterium tuberculosis* dormancy program. J Exp Med 198:705–13.

12. Abramovitch RB, Rohde KH, Hsu FF, Russell DG. 2011. *aprABC*: a *Mycobacterium tuberculosis* complex-specific locus that modulates pH-driven adaptation to the macrophage phagosome. Mol Microbiol 80:678–94.

13. Baker JJ, Johnson BK, Abramovitch RB. 2014. Slow growth of *Mycobacterium tuberculosis* at acidic pH is regulated by *phoPR* and host-associated carbon sources. Mol Microbiol 94:56–69.

14. Leistikow RL, Morton RA, Bartek IL, Frimpong I, Wagner K, Voskuil MI. 2010. The *Mycobacterium tuberculosis* DosR regulon assists in metabolic homeostasis and enables rapid recovery from nonrespiring dormancy. J Bacteriol 192:1662–70.

15. Wayne LG, Hayes LG. 1996. An in vitro model for sequential study of shiftdown of *Mycobacterium tuberculosis* through two stages of nonreplicating persistence. Infect Immun 64:2062–9.

16. MacMicking JD, North RJ, LaCourse R, Mudgett JS, Shah SK, Nathan CF. 1997. Identification of nitric oxide synthase as a protective locus against tuberculosis. Proc Natl Acad Sci U S A 94:5243–8.

17. Nathan C, Shiloh MU. 2000. Reactive oxygen and nitrogen intermediates in the relationship between mammalian hosts and microbial pathogens. Proc Natl Acad Sci U S A 97:8841–8.

18. Weinberg JB. 1998. Nitric oxide production and nitric oxide synthase type 2 expression by human mononuclear phagocytes: a review. Mol Med 4:557–91.

19. Yang CS, Yuk JM, Jo EK. 2009. The role of nitric oxide in mycobacterial infections. Immune Netw 9:46–52.

20. Sherman DR, Voskuil M, Schnappinger D, Liao R, Harrell MI, Schoolnik GK. 2001. Regulation of the *Mycobacterium tuberculosis* hypoxic response gene encoding alpha - crystallin. Proc Natl Acad Sci U S A 98:7534–9.

21. Rodriguez GM, Voskuil MI, Gold B, Schoolnik GK, Smith I. 2002. ideR, An essential gene in *Mycobacterium tuberculosis*: role of IdeR in iron-dependent gene expression, iron metabolism, and oxidative stress response. Infect Immun 70:3371–81.

22. Voskuil MI, Bartek IL, Visconti K, Schoolnik GK. 2011. The response of *Mycobacterium tuberculosis* to reactive oxygen and nitrogen species. Front Microbiol 2:105.

23. Rodriguez GM, Smith I. 2003. Mechanisms of iron regulation in mycobacteria: role in physiology and virulence. Mol Microbiol 47:1485–94.

24. Jones CM, Niederweis M. 2011. *Mycobacterium tuberculosis* can utilize heme as an iron source. J Bacteriol 193:1767–70.

25. Mitra A, Ko YH, Cingolani G, Niederweis M. 2019. Heme and hemoglobin utilization by *Mycobacterium tuberculosis*. Nat Commun 10:4260.

26. Kurthkoti K, Amin H, Marakalala MJ, Ghanny S, Subbian S, Sakatos A, Livny J, Fortune SM, Berney M, Rodriguez GM. 2017. The capacity of *Mycobacterium tuberculosis* to survive iron starvation might enable it to persist in iron-deprived microenvironments of human granulomas. mBio 8:e01092–17.

27. Gold B, Rodriguez GM, Marras SA, Pentecost M, Smith I. 2001. The *Mycobacterium tuberculosis* IdeR is a dual functional regulator that controls transcription of genes involved in iron acquisition, iron storage and survival in macrophages. Mol Microbiol 42:851–65.

28. Pandey R, Rodriguez GM. 2014. IdeR is required for iron homeostasis and virulence in *Mycobacterium tuberculosis*. Mol Microbiol 91:98–109.

29. Pohl E, Holmes RK, Hol WG. 1999. Crystal structure of the iron-dependent regulator (IdeR) from *Mycobacterium tuberculosis* shows both metal binding sites fully occupied. J Mol Biol 285:1145–56.

30. Anand K, Tripathi A, Shukla K, Malhotra N, Jamithireddy AK, Jha RK, Chaudhury SN, Rajmani RS, Ramesh A, Nagaraja V, Gopal B, Nagaraju G, Narain Seshayee AS, Singh A. 2021. *Mycobacterium tuberculosis* SufR responds to nitric oxide via its 4Fe-4S cluster and regulates Fe-S cluster biogenesis for persistence in mice. Redox Biol 46:102062.

31. Kumar A, Toledo JC, Patel RP, Lancaster JR, Jr., Steyn AJ. 2007. *Mycobacterium tuberculosis* DosS is a redox sensor and DosT is a hypoxia sensor. Proc Natl Acad Sci U S A 104:11568–73.

32. Couture M, Yeh SR, Wittenberg BA, Wittenberg JB, Ouellet Y, Rousseau DL, Guertin M. 1999. A cooperative oxygen-binding hemoglobin from *Mycobacterium tuberculosis*. Proc Natl Acad Sci U S A 96:11223–8.

33. Ouellet H, Ouellet Y, Richard C, Labarre M, Wittenberg B, Wittenberg J, Guertin M. 2002. Truncated hemoglobin HbN protects *Mycobacterium bovis* from nitric oxide. Proc Natl Acad Sci U S A 99:5902–7.

34. Pathania R, Navani NK, Gardner AM, Gardner PR, Dikshit KL. 2002. Nitric oxide scavenging and detoxification by the *Mycobacterium tuberculosis* haemoglobin, HbN in *Escherichia coli*. Mol Microbiol 45:1303–14.

35. Blum M, Chang HY, Chuguransky S, Grego T, Kandasaamy S, Mitchell A, Nuka G, Paysan-Lafosse T, Qureshi M, Raj S, Richardson L, Salazar GA, Williams L, Bork P, Bridge A, Gough J, Haft DH, Letunic I, Marchler-Bauer A, Mi H, Natale DA, Necci M, Orengo CA, Pandurangan AP, Rivoire C, Sigrist CJA, Sillitoe I, Thanki N, Thomas PD, Tosatto SCE, Wu CH, Bateman A, Finn RD. 2021. The InterPro protein families and domains database: 20 years on. Nucleic Acids Res 49:D344–d354.

36. Guo Y, Guo G, Mao X, Zhang W, Xiao J, Tong W, Liu T, Xiao B, Liu X, Feng Y, Zou Q. 2008. Functional identification of HugZ, a heme oxygenase from *Helicobacter pylori*. BMC Microbiol 8:226.

37. Cunningham AF, Spreadbury CL. 1998. Mycobacterial stationary phase induced by low oxygen tension: cell wall thickening and localization of the 16-kilodalton alpha-crystallin homolog. J Bacteriol 180:801–8.

38. Rastogi S, Singh AK, Chandra G, Kushwaha P, Pant G, Singh K, Mitra K, Sashidhara KV, Krishnan MY. 2017. The diacylglycerol acyltransferase Rv3371 of *Mycobacterium tuberculosis* is required for growth arrest and involved in stress-induced cell wall alterations. Tuberculosis 104:8–19.

39. Sarathy J, Dartois V, Dick T, Gengenbacher M. 2013. Reduced drug uptake in phenotypically resistant nutrient-starved nonreplicating *Mycobacterium tuberculosis*. Antimicrob Agents Chemother 57:1648–53.

40. Seiler P, Ulrichs T, Bandermann S, Pradl L, Jörg S, Krenn V, Morawietz L, Kaufmann SH, Aichele P. 2003. Cell-wall alterations as an attribute of *Mycobacterium tuberculosis* in latent infection. J Infect Dis 188:1326–31.

41. Pal R, Hameed S, Fatima Z. 2015. Iron deprivation affects drug susceptibilities of Mycobacteria targeting membrane integrity. J Pathog 2015:938523.

42. Rodriguez GM, Sharma N, Biswas A, Sharma N. 2022. The iron response of *Mycobacterium tuberculosis* and its implications for tuberculosis pathogenesis and novel therapeutics. Front Cell Infect Microbiol 12:876667.

43. Grzegorzewicz AE, de Sousa-d’Auria C, McNeil MR, Huc-Claustre E, Jones V, Petit C, Angala SK, Zemanová J, Wang Q, Belardinelli JM, Gao Q, Ishizaki Y, Mikušová K, Brennan PJ, Ronning DR, Chami M, Houssin C, Jackson M. 2016. Assembling of the *Mycobacterium tuberculosis* cell wall core. J Biol Chem 291:18867–79.

44. Hübscher J, Lüthy L, Berger-Bächi B, Stutzmann Meier P. 2008. Phylogenetic distribution and membrane topology of the LytR-CpsA-Psr protein family. BMC Genomics 9:617.

45. Köster S, Upadhyay S, Chandra P, Papavinasasundaram K, Yang G, Hassan A, Grigsby SJ, Mittal E, Park HS, Jones V, Hsu FF, Jackson M, Sassetti CM, Philips JA. 2017. *Mycobacterium tuberculosis* is protected from NADPH oxidase and LC3-associated phagocytosis by the LCP protein CpsA. Proc Natl Acad Sci U S A 114:E8711–e8720.

46. Malm S, Maaß S, Schaible UE, Ehlers S, Niemann S. 2018. *In vivo* virulence of *Mycobacterium tuberculosis* depends on a single homologue of the LytR-CpsA-Psr proteins. Sci Rep 8:3936.

47. Wang Q, Zhu L, Jones V, Wang C, Hua Y, Shi X, Feng X, Jackson M, Niu C, Gao Q. 2015. CpsA, a LytR-CpsA-Psr family protein in *Mycobacterium marinum*, is required for cell wall integrity and virulence. Infect Immun 83:2844–54.

48. Stefanović C, Hager FF, Schäffer C. 2021. LytR-CpsA-Psr glycopolymer transferases: Essential bricks in Gram-positive bacterial cell wall assembly. Int J Mol Sci 22:908.

49. Grosjean N, Yee EF, Kumaran D, Chopra K, Abernathy M, Biswas S, Byrnes J, Kreitler DF, Cheng JF, Ghosh A, Almo SC, Iwai M, Niyogi KK, Pakrasi HB, Sarangi R, van Dam H, Yang L, Blaby IK, Blaby-Haas CE. 2024. A hemoprotein with a zinc-mirror heme site ties heme availability to carbon metabolism in cyanobacteria. Nat Commun 15:3167.

50. Richter AS, Banse C, Grimm B. 2019. The GluTR-binding protein is the heme-binding factor for feedback control of glutamyl-tRNA reductase. Elife 8:e46300.

51. Zhao A, Fang Y, Chen X, Zhao S, Dong W, Lin Y, Gong W, Liu L. 2014. Crystal structure of *Arabidopsis* glutamyl-tRNA reductase in complex with its stimulator protein. Proc Natl Acad Sci U S A 111:6630–5.

52. Aftab H, Donegan RK. 2024. Regulation of heme biosynthesis via the coproporphyrin dependent pathway in bacteria. Front Microbiol 15:1345389.

53. Dailey HA, Gerdes S, Dailey TA, Burch JS, Phillips JD. 2015. Noncanonical coproporphyrin-dependent bacterial heme biosynthesis pathway that does not use protoporphyrin. Proc Natl Acad Sci U S A 112:2210–5.

54. Dailey HA, Dailey TA, Gerdes S, Jahn D, Jahn M, O’Brian MR, Warren MJ. 2017. Prokaryotic heme biosynthesis: Multiple pathways to a common essential product. Microbiol Mol Biol Rev 81:e00048–16.

55. Cheung KM, Spencer P, Timko MP, Shoolingin-Jordan PM. 1997. Characterization of a recombinant pea 5-aminolevulinic acid dehydratase and comparative inhibition studies with the *Escherichia coli* dehydratase. Biochemistry 36:1148–56.

56. Lindblad B, Lindstedt S, Steen G. 1977. On the enzymic defects in hereditary tyrosinemia. Proc Natl Acad Sci U S A 74:4641–5.

57. Donegan RK, Fu Y, Copeland J, Idga S, Brown G, Hale OF, Mitra A, Yang H, Dailey HA, Niederweis M, Jain P, Reddi AR. 2022. Exogenously scavenged and endogenously synthesized heme are differentially utilized by *Mycobacterium tuberculosis*. Microbiol Spectr 10:e0360422.

58. Donegan RK, Moore CM, Hanna DA, Reddi AR. 2019. Handling heme: The mechanisms underlying the movement of heme within and between cells. Free Radic Biol Med 133:88–100.

59. Hanna DA, Harvey RM, Martinez-Guzman O, Yuan X, Chandrasekharan B, Raju G, Outten FW, Hamza I, Reddi AR. 2016. Heme dynamics and trafficking factors revealed by genetically encoded fluorescent heme sensors. Proc Natl Acad Sci U S A 113:7539–44.

60. Santos TMA, Lammers MG, Zhou M, Sparks IL, Rajendran M, Fang D, De Jesus CLY, Carneiro GFR, Cui Q, Weibel DB. 2018. Small molecule chelators reveal that iron starvation inhibits late stages of bacterial cytokinesis. ACS Chem Biol 13:235–246.

61. Ballister ER, Samanovic MI, Darwin KH. 2019. *Mycobacterium tuberculosis* Rv2700 contributes to cell envelope integrity and virulence. J Bacteriol 201:e00228–19.

62. Rath P, Huang C, Wang T, Wang T, Li H, Prados-Rosales R, Elemento O, Casadevall A, Nathan CF. 2013. Genetic regulation of vesiculogenesis and immunomodulation in *Mycobacterium tuberculosis*. Proc Natl Acad Sci U S A 110:E4790–7.

63. Salgueiro-Toledo VC, Bertol J, Gutierrez C, Serrano-Mestre JL, Ferrer-Luzon N, Vázquez-Iniesta L, Palacios A, Pasquina-Lemonche L, Espaillat A, Lerma L, Weinrick B, Lavin JL, Elortza F, Azkargorta M, Prieto A, Buendía-Nacarino P, Luque-García JL, Neyrolles O, Cava F, Hobbs JK, Sanz J, Prados-Rosales R. 2025. Maintenance of cell wall remodeling and vesicle production are connected in Mycobacterium tuberculosis. Elife 13:RP94982.

64. Prados-Rosales R, Weinrick BC, Piqué DG, Jacobs WR, Jr., Casadevall A, Rodriguez GM. 2014. Role for *Mycobacterium tuberculosis* membrane vesicles in iron acquisition. J Bacteriol 196:1250–6.

65. Baumgart M, Schubert K, Bramkamp M, Frunzke J. 2016. Impact of LytR-CpsA-Psr proteins on cell wall biosynthesis in *Corynebacterium glutamicum*. J Bacteriol 198:3045–3059.

66. Over B, Heusser R, McCallum N, Schulthess B, Kupferschmied P, Gaiani JM, Sifri CD, Berger-Bächi B, Stutzmann Meier P. 2011. LytR-CpsA-Psr proteins in *Staphylococcus aureus* display partial functional redundancy and the deletion of all three severely impairs septum placement and cell separation. FEMS Microbiol Lett 320:142–51.

67. Johnsborg O, Håvarstein LS. 2009. Pneumococcal LytR, a protein from the LytR-CpsA-Psr family, is essential for normal septum formation in *Streptococcus pneumoniae*. J Bacteriol 191:5859–64.

68. Gligonov IA, Bagaeva DI, Demina GR, Vostroknutova GN, Vorozhtsov DS, Kaprelyants AS, Savitsky AP, Shleeva MO. 2024. The accumulation of methylated porphyrins in dormant cells of *Mycolicibacterium smegmatis* is accompanied by a decrease in membrane fluidity and an impede of the functioning of the respiratory chain. Biochim Biophys Acta Biomembr 1866:184270.

69. Nikitushkin VD, Shleeva MO, Zinin AI, Trutneva KA, Ostrovsky DN, Kaprelyants AS. 2016. The main pigment of the dormant *Mycobacterium smegmatis* is porphyrin. FEMS Microbiol Lett 363:fnw206.

70. Ganief N, Sjouerman J, Albeldas C, Nakedi KC, Hermann C, Calder B, Blackburn JM, Soares NC. 2018. Associating H_2_O_2_- and NO-related changes in the proteome of *Mycobacterium smegmatis* with enhanced survival in macrophage. Emerg Microbes Infect 7:212.

71. Chouchane S, Girotto S, Kapetanaki S, Schelvis JP, Yu S, Magliozzo RS. 2003. Analysis of heme structural heterogeneity in *Mycobacterium tuberculosis* catalase-peroxidase (KatG). J Biol Chem 278:8154–62.

72. Ouellet H, Lang J, Couture M, Ortiz de Montellano PR. 2009. Reaction of *Mycobacterium tuberculosis* cytochrome P450 enzymes with nitric oxide. Biochemistry 48:863–72.

73. Khare G, Nangpal P, Tyagi AK. 2017. Differential roles of iron storage proteins in maintaining the iron homeostasis in *Mycobacterium tuberculosis*. PLoS One 12:e0169545.

74. Mohanty A, Parida A, Subhadarshanee B, Behera N, Subudhi T, Koochana PK, Behera RK. 2021. Alteration of coaxial heme ligands reveals the role of heme in bacterioferritin from *Mycobacterium tuberculosis*. Inorg Chem 60:16937–16952.

75. D’Aquino JA, Tetenbaum-Novatt J, White A, Berkovitch F, Ringe D. 2005. Mechanism of metal ion activation of the diphtheria toxin repressor DtxR. Proc Natl Acad Sci U S A 102:18408–13.

76. Frunzke J, Gätgens C, Brocker M, Bott M. 2011. Control of heme homeostasis in *Corynebacterium glutamicum* by the two-component system HrrSA. J Bacteriol 193:1212–21.

77. Bibb LA, Kunkle CA, Schmitt MP. 2007. The ChrA-ChrS and HrrA-HrrS signal transduction systems are required for activation of the hmuO promoter and repression of the hemA promoter in *Corynebacterium diphtheriae*. Infect Immun 75:2421–31.

78. Choby JE, Grunenwald CM, Celis AI, Gerdes SY, DuBois JL, Skaar EP. 2018. *Staphylococcus aureus* HemX modulates glutamyl-tRNA reductase abundance to regulate heme biosynthesis. mBio 9:e02287–17.

79. Hanna DA, Hu R, Kim H, Martinez-Guzman O, Torres MP, Reddi AR. 2018. Heme bioavailability and signaling in response to stress in yeast cells. J Biol Chem 293:12378–12393.

80. Rohde KH, Abramovitch RB, Russell DG. 2007. *Mycobacterium tuberculosis* invasion of macrophages: linking bacterial gene expression to environmental cues. Cell Host Microbe 2:352–64.

81. Chen Y, MacGilvary NJ, Tan S. 2024. *Mycobacterium tuberculosis* response to cholesterol is integrated with environmental pH and potassium levels via a lipid metabolism regulator. PLoS Genet 20:e1011143.

82. Chen Y, Hagopian B, Tan S. 2025. Cholesterol metabolism and intrabacterial potassium homeostasis are intrinsically related in *Mycobacterium tuberculosis*. PLoS Pathog 21:e1013207.

83. Kevorkian YL, MacGilvary NJ, Giacalone D, Johnson C, Tan S. 2022. Rv0500A is a transcription factor that links *Mycobacterium tuberculosis* environmental response with division and impacts host colonization. Mol Microbiol 117:1048–1062.

84. Kitzmiller CE, Cheng TY, Prandi J, Sparks IL, Moody DB, Morita YS. 2024. Detergent-induced quantitatively limited formation of diacyl phosphatidylinositol dimannoside in *Mycobacterium smegmatis*. J Lipid Res 65:100533.

